# Interactions Between Dietary Metabolites and Regulatory Risk Variants for Human Colon Cancer

**DOI:** 10.1101/2025.09.05.674475

**Authors:** Tania N. Fabo, Robin M. Meyers, Evin Padhi, Laura N. Kellman, Yang Zhao, Soumya Kundu, David L. Reynolds, Ziwei Chen, Xue Yang, Lisa Ko, Ibtihal Elfaki, Stephen B. Montgomery, Paul A. Khavari

## Abstract

Interactions between genetic variants and environmental factors influence malignancy risk, including for colorectal cancer (CRC). Prevalent CRC susceptibility loci reside predominantly in noncoding regulatory DNA where they may interact with dietary influences to dysregulate expression of specific genes predisposing to neoplasia. The impacts of CRC protective and risk dietary metabolites, butyrate and deoxycholic acid, were thus studied on the transcription-directing activity of 3703 regulatory CRC-associated variants via massively parallel reporter assays (MPRA) in human colonic cells. 1595 variant-dietary metabolite interactions were identified, pointing to dysregulation of MED13L, NKD2, and several modulators of Wnt/β-catenin signaling in potential CRC gene-environment interactions (GxE). Opposing impacts of butyrate and deoxycholic acid were also uncovered, indicating dietary influences may converge on common CRC risk loci and nominating FOSL1 and SP1 as mediators of these opposing responses. Coupling MPRA to relevant environmental factors offers an approach to extend insight into GxE in common human cancers.

## INTRODUCTION

Colorectal cancer (**<u>CRC</u>**) is the second leading cause of cancer death globally^1^. Diet is an important environmental risk factor for CRC, contributing to 30-50% of cases^2,3^. Major dietary risks include a high fat intake, processed red meats, low fiber intake, and alcohol, while high fiber diets, fruits, and vegetables are protective against CRC^3,4^. Inherited genetic risk for CRC ranges along a spectrum from autosomal dominant *APC* mutations in familial adenomatous polyposis to polygenic effects from common noncoding variants with lower effect sizes. While genome-wide association studies (**<u>GWAS</u>**) have identified over 100 susceptibility loci for CRC^5^, the additive genetic effects at these loci fall short of CRC’s “true” heritability, as estimated by twin studies^6,7^ and gene-by-environment (**<u>GxE</u>**) effects are one potential explanation for this so-called “missing” heritability^7^. The detection of GxE effects has proven difficult, however, as population studies are typically underpowered to detect significant interactions^8–12^. Focused experimental approaches may help uncover interactions between dietary and genetic risk factors in CRC pathogenesis.

Butyrate and deoxycholic acid (**<u>DCA</u>**) are dietary metabolites that modulate tumor progression and CRC risk^13–15^. Butyrate is a short-chain fatty acid produced by gut microbiota that is increased with higher fiber intake and that inhibits cancer progression^16^. Butyrate promotes apoptosis, inhibits cellular hyperproliferation, decreases inflammation, and inhibits histone deacetylation. The latter role for butyrate has been linked to suppressed expression of oncogenes and upregulation of tumor suppressor genes^17^. DCA, in contrast, is a secondary bile acid increased by fat intake. DCA’s pro-tumorigenic effects may include increasing oxidative stress, inflammation, and cell proliferation in colonic epithelium^18,19^. DCA regulates pathways implicated in colonic cell proliferation, differentiation, and apoptosis, including the Wnt/β-catenin pathway, which plays a role in hereditary and sporadic CRC^20^.

Here we evaluate interactions between CRC-relevant, diet-modulated environmental influences and CRC-associated noncoding risk variants using massively parallel reporter assays (**<u>MPRAs</u>**). MPRAs screen for transcription-altering effects of thousands of barcoded candidate variants and have been broadly used to identify functional noncoding variants at GWAS-nominated risk loci for various traits, including CRC^21–24^. This approach, termed “**Metabolite-MPRA**”, was applied to both primary human colonic epithelial cells as well as transformed colonic carcinoma cells. Complementing the 808 CRC-risk associated regulatory DNA variants displaying differential constitutive activity, metabolite-MPRA uncovered 1,595 CRC risk-linked variants modulated by butyrate and DCA. These variants were used to nominate transcription factors (**<u>TFs</u>**) whose binding and activity may be altered by disease-linked SNVs as well as corresponding target genes in pathways potentially engaged in CRC GxE.

## RESULTS

### Opposite modulation of CRC pathways by Butyrate and DCA

To examine effects of CRC-relevant dietary metabolites on colonic cell genomic expression and accessibility, RNA-Seq and ATAC-Seq were performed on primary human colon and HCT116 cells stimulated with butyrate or DCA (**Fig. 1A**). Differential analysis of stimulated cells compared to control treated cells was performed to nominate differentially expressed genes and accessible regions. Strong clustering of replicates and conditions across all datasets was observed, as well as strong nucleosome phasing and transcription start site (TSS) enrichment in the ATAC-Seq data (**Supplementary Fig. 1A-C)**. Differential expression analysis nominated 11,812 genes upregulated and 9,290 genes downregulated by butyrate, along with 8,119 genes upregulated and 9,740 genes downregulated by DCA across timepoints and cell types (FDR < 0.05). Differential peak accessibility analysis nominated 77,971 differentially accessible peaks responding to butyrate (representing 61% of total peaks), with 38,480 peaks opening and 39,491 peaks closing (FDR < 0.05; |log2FC| > 1), consistent with prior studies that have found roughly equal number of peaks opening and closing in response to butyrate stimulation, despite its known role as a histone deacetylase inhibitor^25,26^. DCA, a metabolite whose genome regulatory impacts are not well characterized, induced similarly broad changes in accessibility, opening 25,933 regions and closing 33,434 (FDR < 0.05; |log2FC| > 1). These two diet-modified factors therefore broadly altered colonic cell gene expression and genome accessibility.

**Figure 1.**
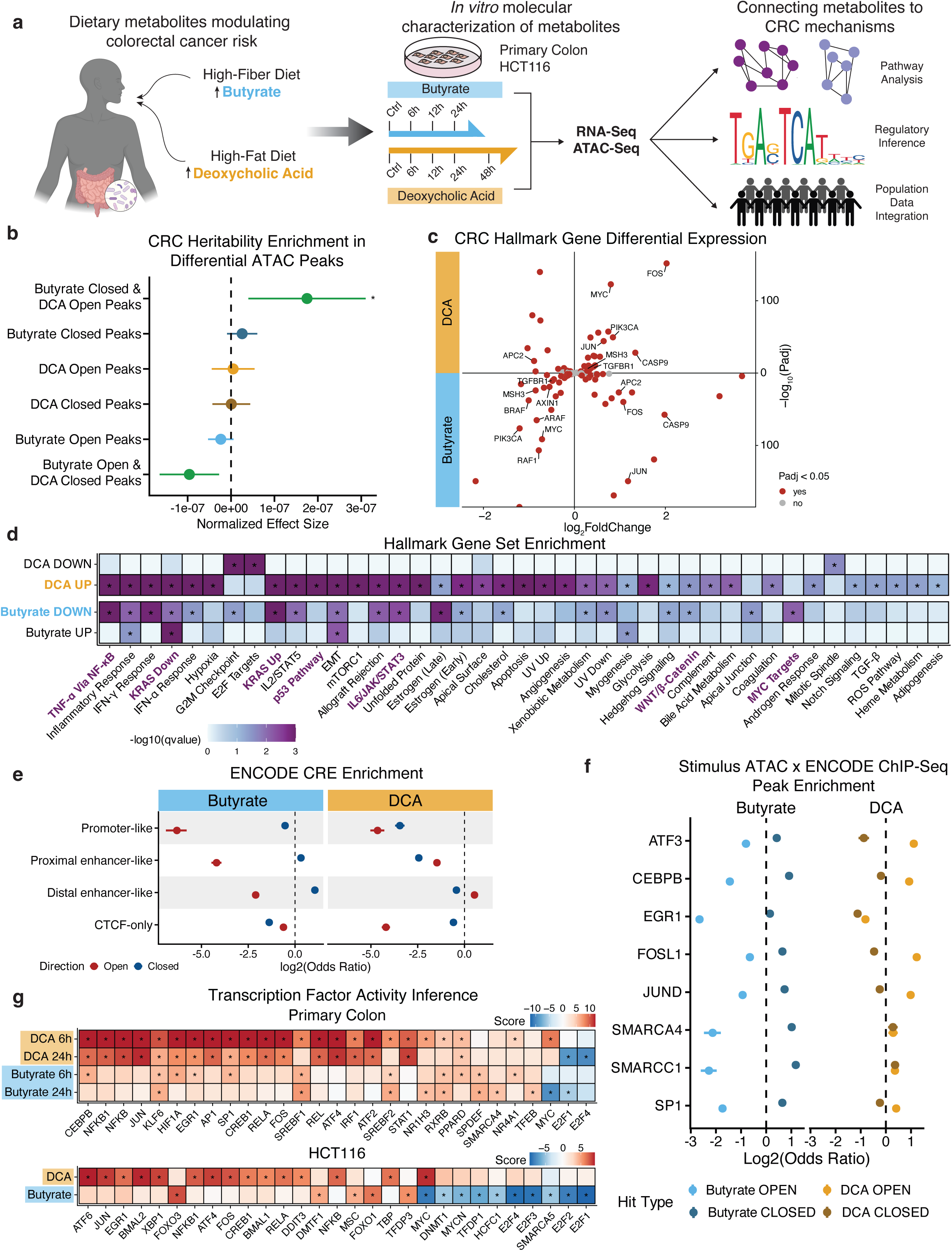
Butyrate and deoxycholic acid as potential modulators of noncoding variant risk in colorectal cancer. (a) Butyrate and DCA RNA-Seq and ATAC-Seq schematic. (b) Partitioned heritability enrichment of differential HCT116 ATAC peaks for colorectal cancer using LD score regression and GWAS summary statistics for CRC; (c) Differential expression of CRC hallmark genes by butyrate and DCA in primary colon cells. (d) Hallmark gene set enrichment analysis for RNA-Seq hit types at 24 hours: butyrate upregulated (UP), butyrate downregulated (DOWN), DCA upregulated (UP), DCA downregulated (DOWN). Genes were filtered by expression (logBM > 1) and ranked by logFC (absolute value). (e) Odds ratio and 95% confidence intervals for enrichment of stimulus-induced differential ATAC peaks in ENCODE cis-regulatory element annotations. (f) Odds ratios and 95% Cl for enrichment of 41 ENCODE ChIP-Seq peak datasets within stimulus-induced differential ATAC peaks. (g) Inference of transcription factor activity from stimulus-induced differentially expressed genes using decoupleR.

To begin to explore a linkage between studied stimuli and CRC risk loci, partitioned heritability analysis was performed on using stratified LD score regression^27–29^. Notably, peaks opened by DCA and closed by butyrate were significantly enriched in heritability, in a pattern of opposing regulation at CRC risk loci by the two stimuli directionally concordant with their known impacts on CRC risk (**Fig. 1B**). The opposite pattern of depletion was found for peaks that are both opened by butyrate and closed by DCA, while no enrichment was found for overlapping peaks which were opened or closed by both stimuli (**Supplementary Fig. 2A**). A similar analysis performed for top genes modulated by the stimuli found that genes that had a significant difference in their response to butyrate versus DCA (“But v DCA Hits”) were the most strongly enriched for CRC heritability (**Supplementary Fig. 2B**), underscoring the role for butyrate and DCA as opposing stimuli in CRC.

The impacts of butyrate and DCA on CRC-relevant gene expression were next evaluated, beginning with differential expression of hallmark CRC genes. Oncogenes, such as *MYC*, were differentially expressed in opposite directions by the stimuli (**Fig. 1C)**. Gene set enrichment analysis revealed enrichment for both stimulus-specific and shared general and hallmark cancer-relevant gene pathways, such as bile acid metabolism for DCA and negative MYC regulation by butyrate **(Fig. 1D)**. Notably, enrichments validated findings on the opposing effects of butyrate and DCA on carcinogenic transcriptional pathways, such as their respective repression and activation of Wnt/β-catenin^20,30^, TNF-ɑ/NF-κB^31,32^, and IL6/JAK/STAT3 signaling^33,34^, further supporting previously described roles of butyrate as anti-tumorigenic and anti-inflammatory^35–37^ and DCA as pro-tumorigenic and pro-inflammatory^38,39^.

ENCODE annotation enrichment showed that genomic regions opened by butyrate were strongly depleted for promoter and enhancer annotations relative to the closed regions, while these opened peaks were strongly enriched for ChromHMM heterochromatin and quiescent/low transcription (**Fig. 1E; Supplementary Fig. 2C**). These findings are consistent with butyrate’s role as an HDAC inhibitor^40^. Further, ENCODE distal enhancer regions and ChromHMM weak transcription annotations were observed to be enriched in both butyrate closed and DCA open peaks, consistent with a previous study that compared GWAS and eQTL hits and found, among many systemic differences, that GWAS hits tended to be distal to TSSs and immersed in more complex regulatory architecture^41^. To nominate specific TFs mediating opposing regulatory mechanisms at differential peaks, stimulus-modulated ATAC-Seq peaks were intersected with HCT116 TF ChIP-Seq peaks from the ENCODE dataset. The binding of AP-1 TFs, ATF3, FOSL1, and JUND, as well as SP1 and CEBPB was enriched in butyrate closed and DCA open peaks (**Fig. 1F; Supplementary Fig. 2D**). These TFs are known to regulate oncogenic and inflammatory pathways whose activities have been previously shown to be modulated with each stimulus, though not in an opposite manner^42–50^. decoupleR was used to further infer TF activity from differentially expressed genes. This approach provided evidence for butyrate’s negative regulation of *MYC*, as well as potentially novel regulatory interactions, such as butyrate’s upregulation of *FOXO1*, a negative regulator of adipogenesis, and DCA’s activation of CREB1, an oncogenic TF (**Fig. 1G**). These TFs represent candidate regulators at sites of regions of the genome whose accessibility is altered by DCA and butyrate.

### MPRA captures dynamic and opposing regulation by butyrate and DCA at CRC risk loci

To examine butyrate and DCA impacts on transcription-directing activity of CRC risk variants, metabolite-MPRA was performed with both normal human colonic epithelial cells and HCT116 CRC cells. Given the relatively low and high transduction efficiency of primary colon cells and cancer cells, respectively, two lentiviral libraries were constructed to accommodate these differences. For the primary colon cells, a small, “hard” barcoded MPRA library of variants placed in their 155-bp genomic context was designed, with 10 barcodes per allelic fragment (**Supplementary Fig. 3A**). For HCT116, a larger, more complex library was designed with “soft” or random barcoded oligos and increased fragment size (200-bp) and including enhancer and TF motif positive controls (**Fig. 2A; Supplementary Fig. 3B**). CRC-associated tag variants for both libraries were expanded to include all variants in linkage disequilibrium (R^2^ > 0.8) and filtered through colon and CRC epigenomic peaks to enrich for variants in regulatory regions. The final primary colon cell MPRA library consisted of 1,595 variants (3,190 alleles) in the forward orientation; the final HCT116 MPRA library consisted of 3,703 variants (7,406 alleles), tested in both forward and reverse complement orientation (**Supplementary Fig. 3C**).

**Figure 2.**
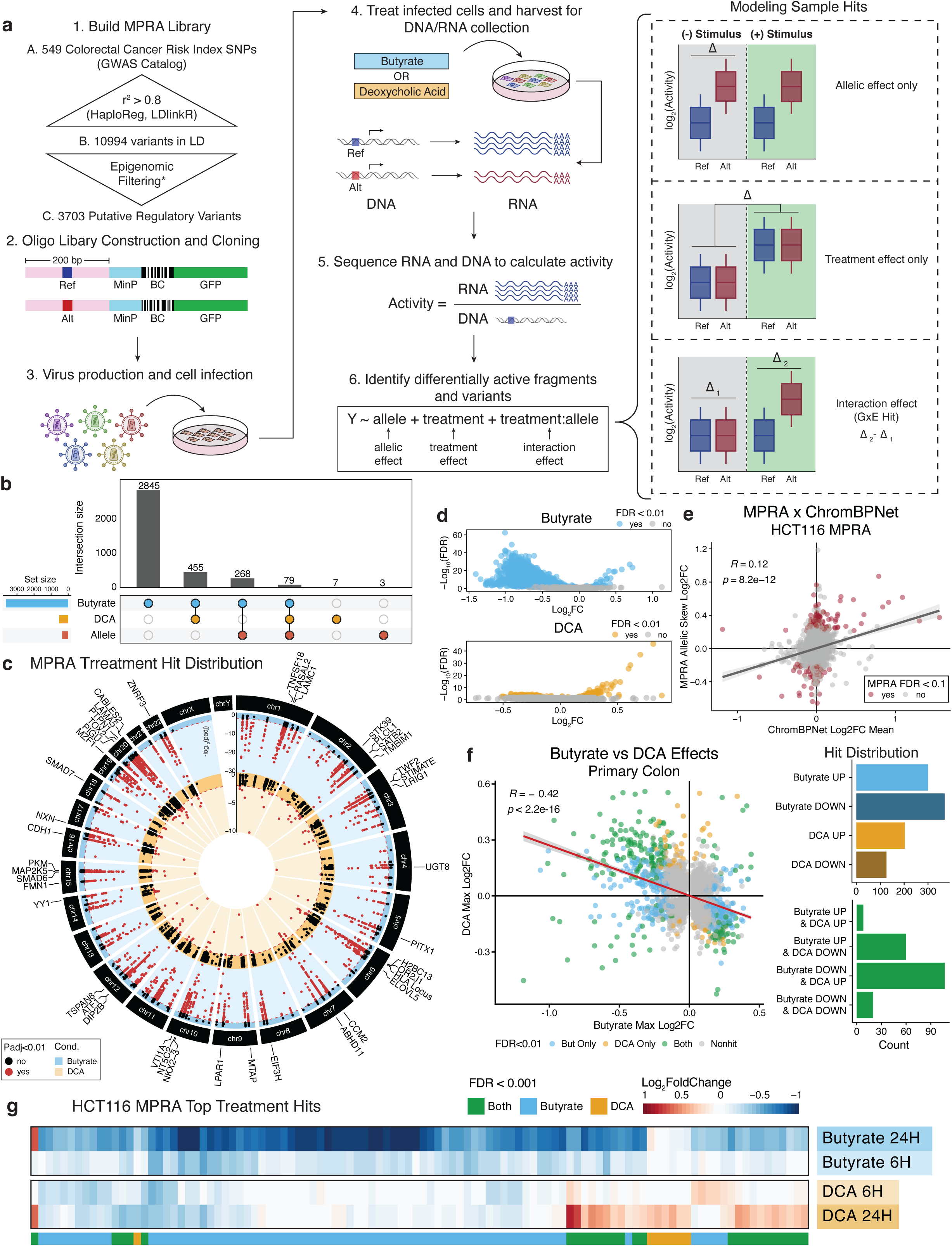
MPRA captures dynamic and opposing regulation by butyrate and DCA at CRC risk loci. (a) HCT116 Metabolite-MPRA schematic. (b Upset plot of HCT116 MPRA treatment and allelic hits. SNPs are called “hits” if treatment FDR< 0.01 for either allele in either orientation and if allelic FDR < 0.1 in either orientation. (c) Circular Manhattan plot showing genome-wide distribution of HCT116 butyrate and DCA treatment hits. (d) Volcano plot of treatment log2FCs and adjusted p-value for butyrate and DCA HCT116 MPRAs. (e) Correlation of MPRA baseline allelic skews and ChromBPNet-predicted mean log2FC variant scores. (f) Correlation of butyrate and DCA treatment log2FCs in primary colon MPRA. (g) Heatmap of treatment and time-dependent effects of butyrate and DCA for top overall treatments hits in HCT116 MPRA (top 50 per condition, Padj < 0.01).

Activity for each fragment with respect to butyrate, DCA, and control was calculated by normalizing RNA counts to DNA counts. For each variant, three types of effects were computed: (1) “allelic” effects, or the difference in activity between reference and alternate alleles; (2) “treatment” effects, or allele-independent response of fragments to either metabolite; and (3) “interaction” effects, or allelic differences that change depending on the condition. The library controls highlighted the MPRA’s ability to read out regulatory effects and perturbations, as controls demonstrated moderate correlation of activities between forward and reverse complement sequences across all fragments (R = 0.49, P < 2.2 x 10^-16^), higher activity of positive controls compared to GWAS fragments and random scrambles, and sensitivity to both motif density and disruption for several active TF motif controls, such as AP-1, NF-κB, and CEBPB (**Supplementary Fig. 3D-E, Supplementary Fig. 4, Supplementary Fig. 5**).

Metabolite-MPRA’s capacity to nominate functional risk variants and the potential mechanisms they engage at baseline through allelic effects and to capture dietary metabolite effects was next assessed. Analysis of metabolite-MPRA identified a total of 458 significant allelic effects (FDR < 0.1), 464 significant butyrate treatment effects and 240 significant DCA treatment effects (FDR < 0.01) in the primary colon MPRA, and 350 allelic effects, 3331 butyrate effects, and 272 DCA effects detected in the HCT116 MPRA (**Fig. 2B-D; Supplementary Fig. 6A-B**). Dietary metabolite effects were not concentrated at any specific genomic locus but were instead broadly distributed across CRC risk loci (**Fig. 2C**). The overwhelming butyrate response in the HCT116 MPRA is hypothesized to be due to the increased levels of non-metabolized butyrate in HCT116 cells caused by the Warburg effect^51^. MPRA-nominated allelic hits represent a set of functional variants at CRC risk loci. The concordance between MPRA allelic skews and variant scores from other genomic assay-based predictive models was first evaluated. Using the ChromBPNet deep learning model trained on HCT116 ATAC peaks, a significant, albeit weak, correlation was observed between HCT116 MPRA allelic effects and ChromBPNet-predicted variant effects on chromatin accessibility (R = 0.12, P = 8.2 x 10^-12^) (**Fig. 2E**). The correlation was driven by a subset (19%) of MPRA hits with a significant ChromBPNet score, while the remaining MPRA hits did not have a predicted effect on chromatin accessibility, suggesting alternate mechanisms of differential risk for these putatively causal variants. Concordance of MPRA allelic hits with variant scores from deltaSVM models trained on SNP-SELEX TF binding data was further evaluated and similarly demonstrated significant global concordance amongst MPRA allelic effects and model-predicted variant scores (Mann Whitney U test P = 0.00151), while concordance was absent in other tested sites (P = 0.423) (**Supplementary Fig. 6C**). These findings suggest differential TF binding and DNA accessibility effects as potential mediators of allelic skew in our MPRAs.

Given the opposing effects of butyrate and DCA at CRC-associated DNA variants, their treatment effects on the same fragment were next compared. In the primary colon MPRA, a moderate negative correlation of treatment effects between the stimuli was observed (ρ = -0.41, P = 1.01 x 10^-179^), with the greatest overlap amongst hits (56.4%) being between variants whose activity is decreased by butyrate (butyrate down) and increased by DCA (DCA up) (**Fig. 2F**). While the same global negative correlation of treatment effects was not observed in the HCT116 MPRA, a sizable decrease in activity in response to butyrate was observed, with a few exceptions, suggesting an inhibitory response to butyrate at CRC risk loci (**Fig. 2G; Supplementary Fig. 6D)**. Comparing treatment effects across cell types, shared treatment effects were positively correlated, strongly for DCA (R = 0.55, P < 2.2 x 10^-16^) and more weakly for butyrate effects (R = 0.18, P = 7.6 x 10^-05^) (**Supplementary Fig. 7**). Taken together, these findings support heritability enrichment analyses and further indicate that these metabolites may have opposite effects at CRC risk loci, with a directionality aligned with their overall effects on risk for CRC.

### Decoding MPRA treatment effects nominates stimulus-responsive TFs in CRC risk

To explore regulatory mechanisms mediating the observed allelic and treatment effects, the location of these hits, stratified by direction of effect, was assessed in the regulatory annotations used to filter the libraries. This focused primarily on the HCT116 MPRA, with similar trends shown in the primary colon data (**Supplementary Table 6-7**). DCA hits were split into up (n=145) and down (n=399) effects. Given the large number of butyrate effects, a high confidence set of effects was nominated by thresholding all fragments on the 90th percentile of the average activity of scramble negative controls in baseline and butyrate conditions. From there, fragments were separated into the following buckets: butyrate up (n=20), butyrate down (n=397), and active & unchanged (n=129). The final category helped disentangle true stimulus effects from stimulus-independent MPRA activity. Butyrate up and down hits, as well as DCA up hits, were strongly enriched for open chromatin (ATAC, DHS), as well as promoter and enhancer histone marks, while DCA down hits were depleted for accessibility (**Fig. 3A**). These patterns were shared with active unchanged fragments, indicating that studied metabolites act primarily on active fragments, with allelic hits, by contrast, minimally enriched for regulatory annotations.

**Figure 3.**
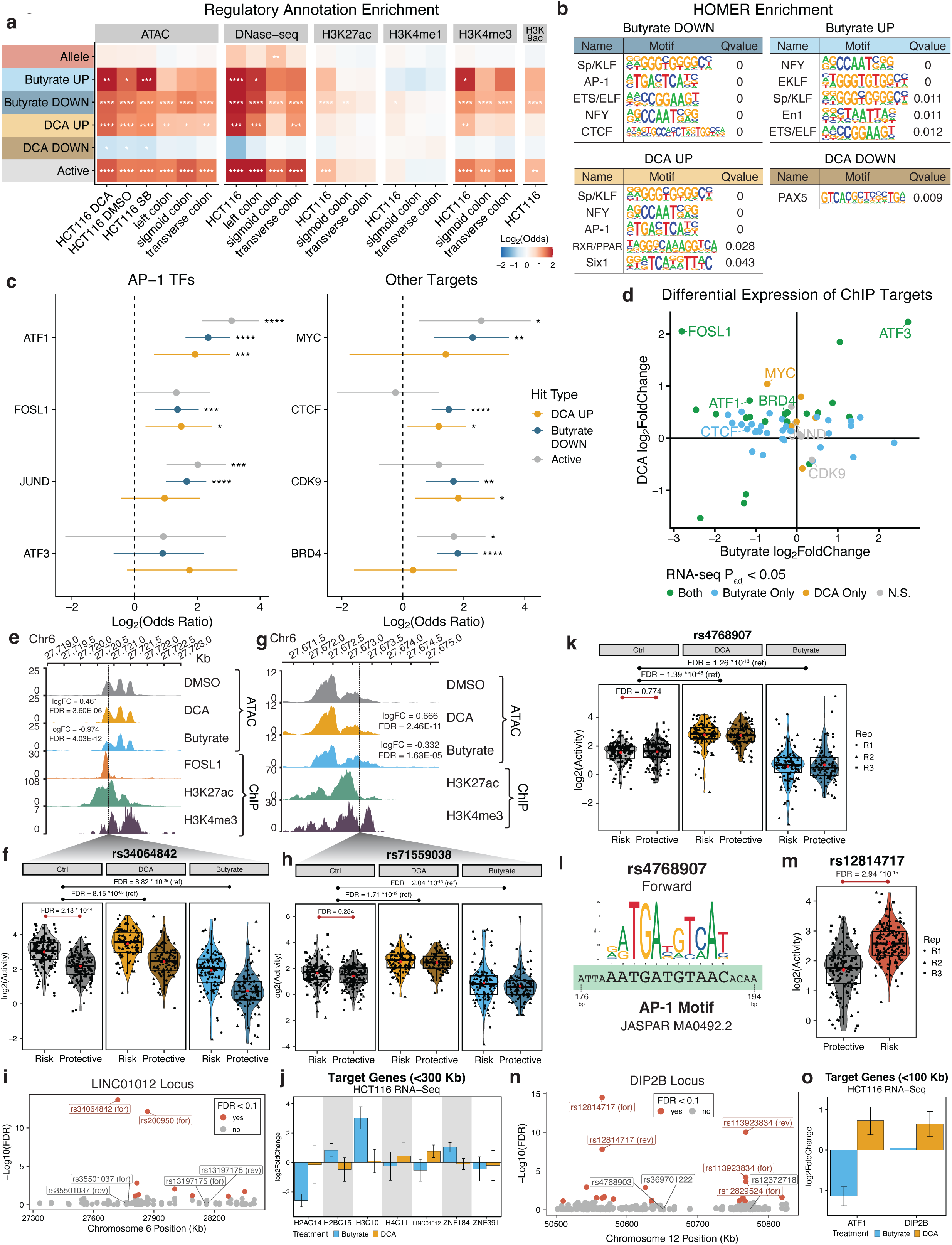
Decoding MPRA treatment effects nominates stimulus-responsive TFs in CRC risk. (a) Enrichment of allelic hits, treatment hits, and active fragments of HCT116 MPRA in colon cell regulatory annotations. (b) HOMER motif enrichment of HCT116 MPRA treatment effects, categorized by treatment and direction. (c) Odds ratios and 95% CI for enrichment of HCT116 MPRA treatment hit overlap with AP-1 TF peaks from HCT116 ENCODE ChIP-seq datasets. (d) Stimulus-induced differential expression in HCT116 cells of all ChIP targets. (e) rs34064842 genome plot. (f) rs34064842 allele HCT116 MPRA activities at baseline and under DCA and butyrate stimulation. (g-h) Same as (e-f) for rs71559038. (i) LINC01012 locus allelic differential activities nominate rs34064842 and rs200950 as causal variants in HCT116 MPRA. (j) Differential expression of genes +/-300 Kb of either variant. (k) rs4768907 allele HCT116 MPRA activities at baseline and under DCA and butyrate stimulation. (l) rs4768907 enhancer fragment contains AP-1 motif. (m) rs12814717 differential activity at baseline in HCT116 MPRA. (n) DIP2B locus allelic differential activities nominate rs12814717 and rs113923834 as causal variants in HCT116 MPRA. (o) Differential expression of genes +/-100 Kb of either variant.

TFs responsible for metabolite effects were next nominated, focusing specifically on butyrate down hits and DCA up hits. HOMER motif analysis enriched for AP-1 and Sp family TFs in butyrate down and DCA up fragments, although the true mediators of these effects remain challenging to decipher, given the high degree of homology between motifs for these TFs (**Fig. 3B**). To determine how MPRA hits overlapped with true TF binding sites, MPRA treatment effects were intersected with ENCODE ChIP-Seq binding peaks for TFs and other regulators. This showed widespread enrichment for butyrate down, DCA up, and active unchanged fragments, with a few notable exceptions (**Supplementary Fig. 8A**). Given the enrichment of AP-1 TF motifs in treatment hits, specific TFs within this family were assessed for mediating the observed effects. Among AP-1 TFs, FOSL1 was the sole TF enriched for binding at butyrate down and DCA up hits (**Fig. 3C**). Differential expression of these TFs further highlighted FOSL1 downregulation by butyrate and upregulation by DCA (**Fig. 3D**). While RNA levels represent only one potential mechanism by which stimuli may regulate TFs, these observations nominate FOSL1 as an AP-1 TF mediating opposite butyrate and DCA effects at CRC risk loci.

Beyond AP-1 TFs, additional enrichments unique to butyrate down and DCA up hits were observed (**Fig. 3C, Supplementary Fig. 8A**). For example, CTCF and RAD21 binding were enriched in butyrate down and DCA up hits. CTCF and RAD21, the latter being part of the cohesin complex, are important for mediating distal enhancer-promoter looping^52^. CDK9 followed a similar pattern of enrichment and has been shown to serve as a key TF regulating oncogenic pathways in CRC^53^. MYC and BRD4 were most strongly enriched in butyrate down hits, while POU2F1 was uniquely enriched amongst these hits, both supporting MYC and nominating BRD4 and POU2F1 as known cancer regulators that may mediate butyrate effects at CRC risk loci.

Metabolite-MPRA effects impact diverse CRC risk loci. At the LINC01012 locus, two treatment-responsive enhancer variants, rs34064842 and rs71559038, displayed activity increased by DCA and decreased by butyrate (**Fig. 3E-H**). Both variants lie in separate differentially accessible ATAC peaks that are opened by DCA and closed by butyrate, while rs34064842 additionally lies in a FOSL1 binding peak. Furthermore, rs34064842 is one of two putatively causal allelic variants at the locus, with the risk allele having significantly higher activity compared to the protective allele (**Fig. 3F,I**). Both enhancer fragments are located closest to LINC01012 (**Fig. 3J**), a long noncoding RNA whose upregulation has been previously linked to cervical cancer, that is upregulated by DCA and downregulated by butyrate in both primary colon and HCT116 cells.

Additional functional CRC-associated variants and a treatment-responsive enhancer were nominated at the *DIP2B* locus. The rs4768907 enhancer fragment is the top treatment hit for DCA and is additionally downregulated by butyrate (**Fig. 3K**). While no TF binding is nominated by extant ChIP-seq data, the fragment is predicted by FIMO to contain an AP-1 binding motif (**Fig. 3L**). The locus also contains several variants with strong allelic skew, including rs12814717, a top allelic hit in both HCT116 and primary colon MPRAs, where the risk allele is more active than the protective allele (**Fig. 3M-N, Supplementary Fig. 8B**). Predicted target genes for variants at this locus include *DIP2B*, the nearest gene and previously shown to promote tumor suppression, and *ATF1*, an AP-1 TF for which rs12814717 is an eQTL in GTEx colon tissue, with the risk variant associated with higher ATF1 expression (**Supplementary Fig. 8C**). Both genes are upregulated by DCA, with *ATF1* downregulated by butyrate (**Fig. 3O)**. These findings highlight how MPRA can be used to nominate both causal variants across loci and enhancers responding to stimuli implicated in CRC risk.

### Identification and characterization of stimulus-modulated CRC risk variants

MPRA identified CRC variants with metabolite-dependent interaction effects, i.e. GxE variants whose allelic differential activity was modulated by either (or both) stimuli. The MPRAs identified 80 butyrate and 17 DCA significant GxE effects in primary colon and 1374 butyrate and 124 DCA significant effects in HCT116 cells (FDR <0.1) (**Fig. 4A-B**). While there were no overlapping GxE hits across MPRAs, several top interaction hits trended in similar directions. These findings highlighted the ability of MPRA to nominate GxE effects for further study that may not be powered for detection in GWAS.

**Figure 4.**
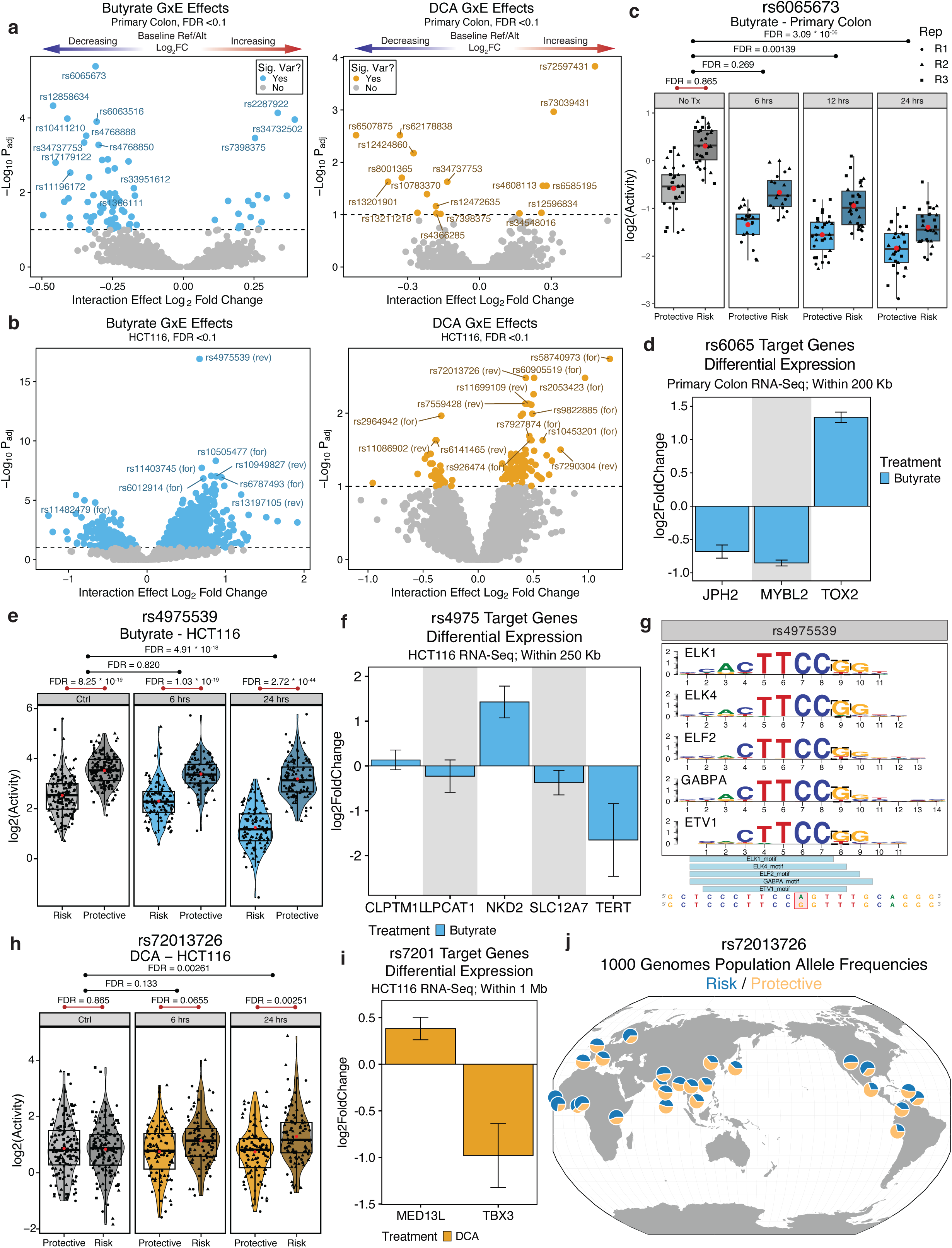
MPRA nominates novel GxE effects at CRC risk loci. (a) Volcano plot of butyrate and DCA interaction (GxE) effects in primary colon MPRA. (b) Same as (a) for HCT116 MPRA. (c) rs6065673 is a butyrate interaction hit in primary colon MPRA. (d) Differential expression of putative target genes for rs6065673 within +/-200 Kb. (e) rs4975539 is a butyrate interaction hit in HCT116 MPRA. (f) Differential expression of putative target genes for rs4975539 within +/-250 Kb. (g) motifbreakR-predicted disruption of TF motifs at rs4975539. (h) rs72013726 is a butyrate interaction hit in primary colon MPRA. (i) Differential expression of putative target genes for rs72013726 within +/-1 Mb. (j) Population-wide distribution of rs72013726 alleles from 1000 Genomes.

At the *TOX2* locus, the primary colon MPRA nominated rs6065673 as a top GxE hit. The variant is a top putative causal variant at the locus at baseline, with the risk allele being more active than the protective allele (**Fig. 4C**). Butyrate treatment both decreases the activity of both fragments and decreases the allelic skew, thereby decreasing the risk gap between alleles. rs6065673 is an eQTL for *TOX2*, whose expression is increased by butyrate and has been shown to function as both a tumor promoter and suppressor, although not in CRC (**Fig. 4D**). Two additional nearby expressed genes include *JPH2*, a cardiomyocyte gene with no established links to cancer, and *MYBL2*, or B-Myb, a proto-oncogene shown to promote progression in CRC and downregulated by butyrate. An alternative example of a butyrate GxE effect was rs4975539, which emerged as a top GxE hit in both forward and reverse complement orientations. rs4975539 is the top allelic hit at the TERT locus at baseline, with the protective allele more active than the risk allele (**Fig. 4E**). Butyrate stimulation, in this case, preferentially decreases activity of the risk allele. While neither rs4975539 nor its tag SNP rs6875445 lie in an eQTL, their nearest gene is *TERT*, a well-established promoter of tumor progression whose expression is decreased by butyrate (**Fig. 4F**). However, the high expression of the protective allele relative to risk make an oncogenic target for this variant less probable. By contrast, *NKD2* is the second closest differentially expressed gene, upregulated by butyrate, and has tumor suppressive functions as an inhibitor of WNT signaling. rs4975539 is predicted to disrupt the binding of several ETS family TFs (**Fig. 4G**), including GABPA, which has been previously shown to synergize with Sp1 to activate transcriptional activity in cancer^54^.

DCA interaction effects further illustrated how stimuli may highlight allelic differences. The *MED13L*-*TBX3* locus was linked to CRC risk in 3 GWAS in European and Asian populations^5,55,56^. At baseline, there was no variant at the locus with allelic skew (**Supplementary Fig. 9A**). rs72013726 emerged as a differentially active variant, however, upon stimulation, with the risk allele being preferentially activated by DCA (**Fig. 4H**). There are only two genes within 1 Mb of this locus, *MED13L* and *TBX3*, both distal from the variant (**Fig. 4I**). MED13L has been implicated in promoting malignant phenotypes in lung cancer^57^, while TBX3 has context-dependent behavior as a either a tumor suppressor or oncoprotein^58,59^. DCA increases MED13L expression and decreasing TBX3 expression, implicating the potential function of these proteins in diet-modulated CRC. Furthermore, rs72013726 exhibited population-wide differences in allele frequency (**Fig. 4J**). While the global, and European, allele frequency was roughly split between risk and protective alleles, the latter was significantly overrepresented in East Asian populations (64.3%; Chi-square p < 10^-5^), while the risk allele was conversely overrepresented in African ancestry populations (64.0%; Chi-square p = 1.1 * 10^-5^). These findings illustrate how gene-environment risk may be further stratified across populations.

A global view at MPRA variants and their putative target genes via PPI network construction revealed a complex network centered on key proteins like MYC and GAPDH, connected to pathways like β-catenin regulation and SMAD signaling (**Supplementary Fig. 10**). The hit-annotated network further displays a high degree of regulatory effects by butyrate and DCA at elements linked to these genes, in both allele-dependent and independent manners. Putative target genes of CRC risk variants modulated by dietary metabolites thus point to pathways implicated in CRC pathogenesis.

## DISCUSSION

Diet plays an important role in stratifying CRC risk and there is a need to understand how dietary stimuli modulate CRC-relevant pathways, including at noncoding regulatory loci harboring risk variants. Here, we profile transcriptomic and genomic accessibility in response to two dietary microbiome metabolites, butyrate and DCA, to characterize their CRC-relevant impacts on 3703 CRC-associated regulatory variants to identify 1595 variant-dietary metabolite interactions. Partitioned heritability analysis uncovered a unique enrichment for differentially accessible peaks both opened by DCA and closed by butyrate at CRC risk loci, providing support for metabolite-modulated variant effects as potentially reflecting putative GxE influences at these sites. Dysregulation of MED13L, NKD2, and Wnt/β-catenin modulators were identified as potential CRC GxE. Opposing impacts of butyrate and deoxycholic acid suggest that dietary influences may converge on loci common that mediate CRC risk.

MPRAs using libraries tailored to differing transduction efficiencies of primary colonic epithelial cells and HCT116 CRC cells showed a strong butyrate down responses, particularly in the HCT116 MPRA, and more modest DCA up responses, with the treatment effects being moderately anticorrelated in the primary colon MPRA. The complex response captures the larger effects of non-metabolized butyrate in HCT116 cells and overall mimics the directionality of the opposing ATAC peaks enriched for CRC risk. Sp1 and FOSL1 were nominated as a TFs that may mediate opposing stimulus responses at putative enhancers, with further analysis showing that butyrate down and DCA up hits are uniquely enriched for distal promoter-enhancer looping, a process which both TFs have been implicated in regulating^60–62^. It is noted that TF peak overlaps were performed based on ChIP-seq experiments performed in baseline conditions. In the future, it will be important to determine how the localization of nominated stimulus-responsive TFs and regulators changes at responsive sites. Importantly, the identification of stimulus-responsive enhancers can still nominate key sites of allele-agnostic impacts of the environment on expression on CRC risk genes, as seen for example with the strong treatment-responsive enhancer at rs4768907 near *ATF1* and *DIP2B*.

Specific variants exhibited potential GxE effects, whereby metabolites modulated variant allelic effects. These variants highlight how environmental stimuli may converge on common risk variants to reduce, enhance, or unmask allelic differences mediating mechanisms of cancer risk. At the *TOX2* locus, the butyrate interaction hit rs6065673 was identified, whose baseline allelic difference is decreased by butyrate; integrating differential expression and eQTL data nominated *MYBL2* and *TOX2* as putative target genes. At the *TBX3* locus, rs72013726 showed how DCA unmasked a putatively causal variant, located distal to putative target genes, *MED13L* and *TBX3*. Global target gene analysis illuminated the importance of stimulus-modulated driver proteins like MYC, and associated networks such as Wnt/β-catenin signaling. Definitive nomination of target genes from MPRA hits, however, remains an ongoing challenge. While target genes were predicted by integrating data from a combination of sources, including eQTL databases, HiChIP looping, and stimulus-induced differential expression, performing perturbation-based assays like CRISPRi will be required to further enhance the accuracy of regulatory variant-target gene assignments. The rs72013726 locus further illustrates how variant risk can further stratify across populations due to differences in allele frequencies, highlighting the importance of including global populations in genomics research.

An ongoing challenge in functional genomics is integration of orthogonal methods of causal variant nomination, an effort further complicated by GxE effects and the limited prediction tools that exist for these. While baseline allelic hits correlate modestly with computational prediction tools like ChromBPNet, an important future step will be to determine how stimulus-informed models can predict interaction effects. More generally, MPRA, as an artificial system, comes with a set of technical limitations imposed by assay constraints, from studied fragment length, to cells lines used as well as to conditions and timepoints tested. It is impossible to fully capture the true nature of environmental regulation at risk loci that occurs over a lifetime to drive CRC, underscoring the importance of future work exploring how well MPRA hits validate in population data. Despite these limitations, our efforts demonstrate how MPRA can capture stimulus-induced regulatory effects and nominate novel interaction effects at CRC risk loci.

## Acknowledgments

We thank G. Rayant and K. Fields for their generous support. We would also like to thank Stanford University and the Stanford Research Computing Center for use of the SCG Informatics Cluster, and the Stanford Genomics facility for sequencing. This work was supported by USVA Office of Research and Development and by NIAMS/NIH AR076965 and AR43799 to PAK, by NCI/NIH CA142635 to PAK, by the Atlas of Regulatory Variants in Disease (ARVID) project from NHGRI/NIH U24HG010856 to PAK.

## Author contributions

T.N.F. and P.A.K. conceptualized the project. T.N.F., R.M.M., L.N.M., and Y.Z. contributed to design of the MPRA. T.N.F. and L.K. performed MPRA experiments. T.N.F., R.M.M., and E.P. analyzed the data. T.N.F., R.M.M., S.K., and Z.C. performed supportive analyses. T.N.F., R.M.M., D.L.R., E.P., S.M., and P.A.K. guided methodology development, experiments and data analysis. T.N.F. and P.A.K. wrote the manuscript with input from all authors.

## FIGURE LEGENDS

**Supplementary Figure 1.** (a) PCA plot for primary colon stimulus RNA-Seq. (b) PCA plot for HCT116 stimulus (24 hour) ATAC-Seq. (c) Nucleosome phasing and TSS enrichment of HCT116 stimulus ATAC-Seq.

**Supplementary Figure 2.** (a) LDSC enrichment of peaks closed by both butyrate and DCA and peaks opened by both butyrate and DCA. (b) RNA-Seq LDSC enrichment; butyrate v DCA hits are genes differentially expressed between butyrate and DCA. (c) ChromHMM enrichment of differential ATAC peaks. (d) Complete HCT116 stimulus ATAC x HCT116 ENCODE ChIP peak enrichment.

**Supplementary Figure 3.** (a) Primary colon MPRA library filtering and design and barcode representation in RNA libraries. (b) Design of HCT116 MPRA positive control enhancer windows and TF motif repeats. (c) Median barcodes per fragment in HCT116 MPRA. (d) Activities of forward vs reverse complement GWAS fragments. (e) Activities of TF motif repeats, positive control fragments, GWAS fragments, and scrambles in HCT116 MPRA.

**Supplementary Figure 4.** (a) Pairwise scatter correlation matrix of HCT116 MPRA replicates across conditions. (b) PCA of oligo-level HCT116 MPRA activity across conditions. (c) Activities of positive control fragment windows across conditions.

**Supplementary Figure 5.** Activities of TF motif repeat fragments and mutants in baseline, DCA, and butyrate HCT116 MPRAs.

**Supplementary Figure 6.** (a) Volcano plots of primary colon butyrate and DCA MPRA treatment effects. (b) Volcano plot of HCT116 MPRA allelic effects at baseline. (c) Concordance of primary colon MPRA allelic effects with SNP-SELEX deltaSVM scores for hits and nonhits. (d) HCT116 MPRA MA plots.

**Supplementary Figure 7.** Correlation of primary colon and HCT116 MPRA treatment effect for shared versus non-shared hits.

**Supplementary Figure 8.** (a) Enrichment log2 odds ratio of MPRA hits across all HCT116 ChIP peaks. (b) Primary colon MPRA plot for rs12814717 baseline differential activity. (c) rs12814717 is an ATF1 eQTL in GTEx sigmoid colon tissue.

**Supplementary Figure 9.** (a) Baseline versus DCA treatment allelic effects at MED13L/TBX3 locus nominates rs72013726 as a condition-specific causal variant. (b) rs10411210 is a GxE hit in butyrate primary colon MPRA. (c) rs4608113 is a GxE hit in DCA primary colon MPRA. (d) rs72597431 is a GxE hit in DCA primary colon MPRA. (e) rs72597431 alt allele creates ATF1 motif. (f) Differential expression of rs72597431 target genes.

**Supplementary Figure 10.** (a) Protein-protein interaction (PPI) network of colon cancer linked SNV genes, from colon epithelial cells across two conditions, consisting of 213 nodes (proteins) and 373 edges (interactions). The highlighted genes are part of modules identified by Markov clustering and enriched in GO: Biological Processes with FDR < 0.05. (b) PPI network of colon cancer linked SNV genes, from HCT 116 cells across two conditions, consisting of 499 nodes and 1482 edges. The highlighted genes are part of modules identified by Markov clustering and enriched in GO: Biological Processes with FDR < 0.05.

## Supplementary Data

**Supplementary Data S1:** Overview of paper datasets (MPRA, ATAC, RNA) and external datasets used for analysis

**Supplementary Data S2:** List of MPRA index SNPs, their location, and original study information and population.

**Supplementary Data S3:** List of all MPRA SNPs, including alleles, index SNP, which MPRA(s) it was tested in, and epigenome annotation.

**Supplementary Data S4:** Complete colon MPRA results – logFC, standard error, pvalue, and FDR for allelic, treatment, and interaction effects.

**Supplementary Data S5:** Complete HCT116 MPRA results – logFC, standard error, pvalue, and FDR for allelic, treatment, and interaction effects.

**Supplementary Data S6**: Complete ChromBPNet results (HCT116 control)

**Supplementary Data S7:** HOMER enrichment results by MPRA treatment hit type (butyrate up, butyrate down, DCA up, DCA down) for HCT116 and primary colon MPRAs.

**Supplementary Data S8:** ChIP enrichment results by MPRA treatment hit type for HCT116 and primary colon MPRAs.

**Supplementary Data S9:** ChIP enrichment results by ATAC peak type.

**Supplementary Data S10:** Full LDSC results for ATAC and RNA differential peaks/genes in CRC GWAS.

## METHODS

### Cell culture

Human primary colonic epithelial cells (Cell Biologics, H-6047; Lot #: 041417PNBAA) were cultured in complete epithelial cell medium (Cell Biologics, H-6621). HCT116 cells (ATCC, CCL-247) were cultured in McCoy’s 5A medium (ATCC, 30-2007), supplemented with 10% FBS and 1% P/S.

### Stimulation

Sodium butyrate (Sigma, B5887) was reconstituted in nuclease-free water for a 908 mM stock solution. It was diluted to a working concentration of 2-5 mM in cell culture media for stimulation. Deoxycholic acid (Sigma, D2510) was reconstituted in DMSO for a 50 mM stock solution and diluted to a working concentration of 100-200 uM for stimulation.

### RNA-Seq

Primary colon and HCT116 cells were stimulated in biological triplicates with butyrate or deoxycholic acid, along with the appropriate vehicle control. Cells were stimulated at 50-70% confluency for 6, 12, and 24h for butyrate and 6, 12, 24, and 48h for deoxycholic acid. Total RNA was extracted from HCT116 cells using the RNeasy Plus Mini Kit (Qiagen, 74136). RNA-seq library preparation was performed using the Illumina-compatible QuantSeq 3’ mRNA-Seq Library Prep Kit FWD (Lexogen, 191.96), following the manufacturer’s protocol. The PCR Add-on and Reamplification Kit (Lexogen, 208.96) was used for qPCR to determine optimal PCR cycle number. Final RNA-seq libraries were quantified using the BioAnalyzer High Sensitivity DNA Kit (Agilent, 5067-4626) prior to submission to Novogene for sequencing using 150 base pair paired-end reads on an Illumina NovaSeq 6000 at a depth of 50 million reads per sample. To process the sequencing reads, adapters were trimmed using Cutadapt^63^, followed by alignment to ENSEMBL release 106, genome build GRCh38 using STAR^64^. featureCounts^65^ was used to generate summarized raw gene counts from aligned bam files, which were used as the input for DESeq2 to perform normalization and differential expression analysis^66^.

### RNA-Seq analysis

Significant differentially expressed genes from stimulus RNA-Seq were filtered by adjusted p-value < 0.05, log2FC greater/less than 0, and log10(BaseMean) > 1. We used the clusterProfiler package in R to perform gene set enrichment analysis^67^, using the hallmark gene set signatures from the msigdbr package^68^.

TF regulatory network analysis was performed using the decoupleR package^69^. Using the CollecTRI gene regulatory network and the differentially expressed genes ranked by DESeq2 stat value, decoupleR fit a univariate linear model to calculate regulatory activities for transcription factors in each stimulus condition. The resulting p-values were adjusted for multiple testing using Bonferroni correction.

### ATAC-Seq

HCT116 cells were stimulated in biological triplicates with butyrate or deoxycholic acid, along with the appropriate vehicle control. Cells were stimulated at 50-70% confluency for 6 and 24h. ATAC-seq was performed on 50,000 nuclei using the Omni-ATAC protocol^70^. Briefly, 500,000 viable cells were lysed with digitonin, pelleted, and washed, after which 50,000 nuclei were counted and aliquoted. The <u>Kaestner Lab Omni-ATAC protocol</u> was used for the remainder of the library preparation, including a double-sided bead purification for final library purification. ATAC libraries were submitted to Novogene for paired-end read sequencing on the NovaSeq 6000 at a depth of 50 million reads per sample. ATAC-seq reads were processed using the ENCODE ATAC-seq pipeline (https://github.com/ENCODE-DCC/atac-seq-pipeline). The pipeline includes FASTQ read adaptor trimming (Cutadapt), read alignment (Bowtie2 aligner), post-alignment filtering of mitochondrial reads, unmapped reads, duplicates, and other reads failing QC (SAMtools, MarkDuplicates). BAM files were converted to tagAlign files and MAC2 used for peak calling on pooled and pseudoreplicates^71^. IDR analysis was run on for all replicates to generate a subset of high confidence peaks across replicates.

Read counts per peak were calculated using the bedtools coverage command line utility. IDR optimal peaks were merged across all conditions and tagAlign counts for each condition were used to calculate coverage. Counts were quantile normalized using the preprocessCore package in R, and quantile-normalized counts were used to call differentially accessible peaks using edgeR^72^. Differential peaks were filtered for FDR < 0.05 and |log2FC| > 1.

### MPRA library design

Primary colon MPRA: The colon MPRA library was constructed from 1,595 unique index variants, shown in GWAS to be significantly associated with colorectal cancer (P < 5 x 10^-8^). These index SNPs were expanded in population-matched LD (R2 > 0.8) using HaploReg and LDLinkR to identify all putatively causal variants. We then performed epigenomic filtering using a union set of ENCODE ATAC-Seq, DNAse-Seq, H3K27Ac ChIP-Seq, H3K4me1 ChIP-Seq, H3K4me3 ChIP-Seq, and H3K9Ac ChIP-Seq peaks from primary colon tissue (left colon, sigmoid colon, transverse colon) and colorectal cancer cell lines (HCT116, HT-29, Caco-2, LoVo, SW480 cells). The final library included 1,595 variants centered on their 155-bp genomic context (hg38). Final oligo design included the following, in order: MPRA forward primer binding site, 155-bp CRE sequence, 22-bp random filler, 20-bp barcode, and reverse primer binding site.

HCT116 MPRA: The HCT116 MPRA library was constructed from 549 unique index variants, including SNPs significantly associated with colorectal cancer (P < 5 x 10^-8^) and a set of sub-threshold SNPs which were either 1) identified in populations understudied in colorectal cancer GWAS^73–75^, or 2) nominated for potential GxE interaction effects for colorectal cancer^8^. A full list of index SNPs and the studies and populations in which they were identified can be found in Supp. Table X. These index SNPs were expanded in population-matched LD (R^2^ > 0.8) using HaploReg and LDLinkR to identify all putatively causal variants. We then performed epigenomic filtering using a union set of ENCODE ATAC-Seq, DNAse-Seq, H3K27Ac ChIP-Seq, H3K4me1 ChIP-Seq, H3K4me3 ChIP-Seq, and H3K9Ac ChIP-Seq peaks from primary colon tissue (left colon, sigmoid colon, transverse colon), colorectal cancer cell lines (HCT116, HT-29, Caco-2, LoVo, SW480 cells), and T-reg cells (CD4-positive, CD25-positive, alpha-beta regulatory T cell). We also included stimulus ATAC-Seq peaks in the epigenomic filtering. Indels >5 bp were filtered out. Variants were centered on their 200-bp genomic context (hg38). For indels, in the case of insertions, the alt fragment is exactly 200-bp while the ref fragment is <200; in the case of deletions, the reverse is true, with the ref fragment exactly 200 and the alt fragment <200. Variant fragments were included in both their forward and reverse complement orientation.

The library included several positive controls. We selected 12 top-scoring enhancers from Neumayr et al. 2022, which performed STARR-Seq in HCT116 cells. As these enhancers were 1200-bp in length, 200-bp windows with 100-bp overlaps were tiled across these enhancers, creating 11 200-bp fragments per enhancer. All enhancer fragments were also tested in their reverse complement orientation, resulting in 264 positive control enhancer fragments. Additionally, we selected 12 unique top scoring TF motifs based on the decoupleR regulon inference results for butyrate and deoxycholic acid: JUN/FOS/AP1 (JASPAR <u>MA0099.3</u>, HOCOMOCO <u>JUN.H12CORE.0.P.B</u>), NFKB/RELA/NFKB1 (JASPAR <u>MA0105.2</u>, HOCOMOCO <u>NFKB1.H12CORE.0.PS.A)</u>, ATF2/ATF4 (JASPAR <u>MA0833.1</u>, HOCOMOCO <u>ATF4.H12CORE.0.P.B)</u>, SREBF1/SREBF2 (JASPAR <u>MA0595.1</u>, HOCOMOCO <u>SRBP1.H12CORE.0.P.B</u>), FOXO1/FOXO3 (JASPAR <u>MA0157.2</u>, HOCOMOCO <u>FOXO1.H12CORE.0.PS.A</u>), CEBPB (JASPAR <u>MA0466.1</u>, HOCOMOCO <u>CEBPB.H12CORE.0.P.B</u>), SP1 (JASPAR <u>MA0079.3</u>, HOCOMOCO <u>SP1.H12CORE.0.P.B</u>), TEAD4 (JASPAR <u>MA0809.1</u>, HOCOMOCO <u>TEAD4.H12CORE.0.PS.A</u>), HIF1A (JASPAR <u>MA1106.1</u>, HOCOMOCO <u>HIF1A.H12CORE.0.P.B</u>), MYC (JASPAR <u>MA0147.3</u>, HOCOMOCO <u>MYC.H12CORE.0.P.B</u>), FOXP3 (JASPAR <u>MA0850.1</u>, HOCOMOCO <u>FOXP3.H12CORE.0.P.B</u>), NR1H3-RXRA (JASPAR <u>MA0494.1</u>, HOCOMOCO <u>NR1H3.H12CORE.0.P.B</u>). For each motif, we created a “mutant” form by introducing a point mutation predicted to be disruptive. The motifs (normal or mutant) were inserted into a neutral background template (from Smith et al. 2013, which also did not overlap with open peaks). The reverse complement of the normal and mutated motif was also inserted into the same (forward orientation) template. The result was 192 additional test fragments for TF motifs.

Negative control fragments were designed by taking a random 250-member subset of the library and randomly scrambling the sequences. A single base mutation was inserted at the center to mimic ref/alt pairs. 15bp PCR adapters were added to the 5’ (AGGACCGGATCAACT) and 3’ (CATTGCGTGAACCGA) end of the 200-bp fragments, resulting in 15,768 230-bp oligos which were synthesized by Agilent Technologies.

### MPRA library construction

Colon MPRA: A 31,900-oligo library was ordered from Agilent Technologies and resuspended to 14 ng/uL. The final library was cloned via a two-step process of 1) inserting the oligos into the pGF MPRA vector; and 2) inserting the minimal promoter and luciferase fragment sequence between the test CRE and barcode. The oligo library was amplified by emulsion PCR using the Micellula DNA Emulsion & Purification Kit (EURx, E3600). pGF (Addgene, 174103) and the amplified oligo library were digested EcoRI and BamHI and ligated together using T4 DNA ligase (NEB, M0202). Cloned library was electroporated into MegaX electrocompetent cells (Thermo Fisher, C640003), which were grown in LB liquid culture under ampicillin selection. The round 1 cloned plasmid and donor vector containing the minimal promoter and luciferase fragment were both digested with XhoI and NheI and ligated together using T4 ligase. Cloned library was electroporated into MegaX and grown in LB+ampicillin. The plasmid DNA library was sequenced to assess oligo barcode frequencies (see below).

HCT116 MPRA: The lentiMPRA library was constructed as described in Gordon et al. (2020). Briefly, a 15,768-oligo library was ordered from Agilent Technologies and resuspended to 20 ng/uL. The first round of PCR added the minimal promoter, followed by a second round of PCR to add 15-bp random barcodes downstream of the promoter. pLS-SceI (Addgene, 137725) was digested with AgeI and SbfI and amplified library inserted into the digested vector using NEBuilder HiFi (NEB, E2621). Cloned library was electroporated into NEB 10-beta electrocompetent cells (NEB, C3020K), which were grown on15-cm carbenicillin agar plates for <16h overnight. Plasmid library was recovered from bacteria using Qiagen Maxi Prep (Qiagen, 12965). The final library was prepared for association sequencing by PCR amplification to add P5 and P7 flowcell adapters and a sample index and submitted to Novogene for sequencing with custom primers that read the CRS/insert (reads 1 & 2), barcode (index read 1), and sample index (index read 2).

### MPRA library production

To produce MPRA lentivirus, 3 million Lenti-X 293T cells (Takara, 632180) were seeded in DMEM (10% FBS) in 15-cm plates and incubated for 2 days until 70-80% confluent. LentiX cells were transfected for viral production using the Lenti-Pac™ HIV Expression Packaging Kit (GeneCopoeia, LT001). MPRA library plasmid was co-transfected with HIV packaging plasmid mix using the Endofectin transfection reagent and incubated for 8h, at which point media was replaced with low serum (2%) DMEM and TiterBoost reagent. Cells were assessed for GFP expression and culture medium collected at 48h post-transfection. Viral supernatant was 50x concentrated using Lenti-X concentrator (Takara, 631232). HCT116 (and primary colon / Jurkat) cells were titrated with lentivirus and MOI calculated as described in Gordon et al. (2020).

### MPRA library infection, DNA/RNA collection, and sequencing library generation

Primary colon cells were transduced with lentivirus at optimal MOI, as calculated by titration. After X days, transduced cells were selected with puromycin for X days. Infected cells were then stimulated with butyrate for 6h, 12h, and 24h or DCA for 12h, 24h, and 48h, or the appropriate vehicle control. After stimulation, cells were lysed in RLT buffer (QIAGEN, 1053393) and stored at -80 C. Library prep was performed as described in [Kellman et al. (2025)]. Briefly, genomic DNA and RNA were isolated from lysed MPRA samples using the AllPrep DNA/RNA Kit (QIAGEN, 80204). mRNA was then isolated from the total RNA using the Dynabeads mRNA DIRECT purification kit (Thermo Fisher, 61011). cDNA synthesis from mRNA samples was performed with SuperScript IV Reverse Transcriptase (Thermo Fisher, 18090010). PCR amplification of cDNA and genomic DNA using PrimeStar Max DNA Polymerase (Takara Bio, R045) added UMIs, sample indexes, and P5/P7 adapters. Pooled libraries were submitted to Novogene for 150bp paired-end sequencing.

HCT116 cells were transduced with library lentivirus at optimal MOI, as calculated by titration. After 2d, transduction efficiency was assessed by checking cells for GFP expression. Infected cells were stimulated with butyrate or deoxycholic acid for 6h and 24h, or the appropriate vehicle control. After stimulation, cells were lysed in RLT buffer (QIAGEN, 1053393) and stored at -80 C. Library prep was performed as described in Gordon et al. (2020). Briefly, genomic DNA and RNA were isolated from lysed MPRA samples using the AllPrep DNA/RNA Kit (QIAGEN, 80204). RNA samples were treated with DNase using the TURBO DNA-free kit (Thermo Fisher, AM1907). cDNA synthesis from RNA samples was performed with SuperScript IV Reverse Transcriptase (Thermo Fisher, 18090010). PCR amplification of cDNA and genomic DNA using using NEBNext Ultra II Q5 Master Mix (NEB, M0544) added UMIs, sample indexes, and P5/P7 adapters. Pooled libraries were submitted to Novogene for sequencing with custom primers that read the barcode (reads 1 & 2), UMI (index read 1), and sample index (index read 2).

### MPRA analysis

Colon MPRA: Barcode and UMI sequences were extracted from FASTQ files for the plasmid DNA library and RNA sample libraries. Barcode sequences were mapped to the barcode reference library using bowtie. PCR duplicates were removed using UMI-tools and summarized barcode counts were subsequently generated. DNA and RNA barcode counts tables were used as input for MPRAnalyze to calculate fragment-level activities and different effect types (allele, treatment, and interaction). MPRAnalyze’s analyzeComparative function was used to calculate normalized effect sizes using likelihood ratio tests, with dnaDesign = ∼ barcode and rnaDesign as follows (full vs reduced): Allele = ∼ allele vs ∼ 1; Treatment = ∼ treatment vs ∼ 1; Interaction = ∼ allele + treatment + allele:treatment vs ∼ allele + treatment. A random effects term added to account for high correlation of barcode activities across replicates.

HCT116 MPRA: Barcodes were mapped to parent oligo sequences by using FLASH to merge paired-end reads derived from the merged sequencing outputs. Random barcodes were extracted from each merged fragment and appended to the read name. STAR v2.7.1a was used to align merged reads against a reference index created from the designed library sequences. After filtering reads that did not map uniquely to a designed sequence or which had low quality alignment scores (< 100), the resulting barcode-oligo pairs were extracted and any sequences detected on multiple oligos were removed. On average, each oligo sequence was mapped to 456 unique barcodes. The MPRAflow count pipeline (https://github.com/shendurelab/MPRAflow) was used to process MPRA sequencing reads, align them to the association dictionary, and generate DNA and RNA barcode counts tables for MPRAnalyze. Given the large number of barcodes (∼500/fragment), a random set of 50 barcodes per fragment, where the sum of pDNA counts (rep1 + 2) was greater than 10, were selected for further analysis. Similar to the colon library analysis, MPRAnalyze was used to normalize RNA counts to DNA counts and perform LRTs to calculate allele, treatment, and interaction effect sizes.

### Enrichment analysis - ChIP-seq datasets

HCT116 ChIP-Seq peaks were downloaded from ENCODE (38 targets) and ReMAP 2022 (40 targets). Additional ChIP-Seq datasets for HDAC1-3 in K562 cells were downloaded from ENCODE. Odds ratios for enrichment of differential ATAC peaks and MPRA treatment hits (FDR < 0.01) in ChIP-Seq peaks were calculated by Fisher’s exact test and adjusted for multiple testing by FDR correction.

### Enrichment analysis - ENCODE CRE

ENCODE candidate cis-regulatory elements were downloaded from SCREEN (https://screen.encodeproject.org/) for HCT116 cells, sigmoid colon, and transverse colon and categorized into promoter-like signatures (PLS), proximal enhancer-like signatures (pELS), distal enhancer-like signatures (dELS), and CTCF-only. Odds ratios for enrichment of MPRA treatment hits (FDR < 0.01) in CRE element categories were calculated by Fisher’s exact test and adjusted for multiple testing by FDR correction.

### Enrichment analysis - ChromHMM

ChromHMM annotations (core 15-state model) for sigmoid colon (E106) were downloaded from the Roadmap Epigenomics Project. Odds ratios for enrichment of MPRA treatment hits (FDR < 0.01) in each chromatin state were calculated by Fisher’s exact test and adjusted for multiple testing by FDR correction.

### HOMER motif analysis

HOMER software for motif discovery and analysis was used to identify motifs enriched in MPRA treatment hits (FDR < 0.01). Figure 3b displays the top known results from the HOMER findMotifsGenome.pl command with default parameters. Full HOMER results can be found in Supp. Table X.

### TF binding disruption prediction

MotifbreakR^76^ was used to predict disruption of transcription factor binding motifs by variants, using HOCOMOCO v11 motif PWMs, using default parameters with a binding threshold of P < 1 x 10^-4^ for at least one allele.

Ren Lab deltaSVM models for 94 TFs based on HT-SELEX data were used to predict differential TF binding at SNPs. The predictions were performed using the Ren Lab <u>web</u> <u>interface</u>. The TF with the maximum absolute deltaSVM score was selected for each SNP for further analysis. Significant MPRA allelic effect variants were those with FDR < 0.1 and alpha/activity > -0.8, while non-significant variants were those with FDR > 0.6 and alpha/activity < -1.0. Differences in deltaSVM distribution between positive and negative allelic effect MPRA variants were calculated by Mann-Whitney U Test.

### Target gene nomination (E2G, eQTLs, HiChIP)

ENCODE-rE2G annotations^77^ for 5 colon sample types (large intestine, left colon, sigmoid colon, transverse colon, and HCT116 cells) were downloaded from ENCODE and MPRA variants were matched to genes based on enhancer/promoter overlap.

Significant eQTL variant-gene pairs were downloaded from the GTEx Portal^78^ (v10, accessed 12/28/24) and eQTLGen^79^ and matched to MPRA variants. GTEx colon samples included sigmoid and transverse colon tissue.

H3K27Ac HiChIP data for colonic epithelial cells from Donohue et al. (2022) was processed as described. Briefly, HiChIP paired-end reads were aligned to hg19 using HiC-Pro and filtered reads processed using hichipper. FitHiChIP was used to call significant chromatin contacts using the default settings except for the following: MappSize=500, IntType=3, BINSIZE=5000, QVALUE=0.01, UseP2PBackgrnd=0, Draw=1, TimeProf=1. HiChIP anchors were mapped to gene transcription start sites using the GENCODE v47 gene annotations. Loop anchors were intersected with MPRA fragments and TSSs independently using bedtools pairtobed. SNP-gene pairs were called if one anchor contained a GWAS SNP and the opposing anchor contained a TSS.

### Misc. Visualization

Genome track plots (Figure 3e,g) were built with pyGenomeTracks^80,81^. Circos Manhattan plot of MPRA variants (Figure 2c) was built with the circlize package in R^82^.

### PPI Network

Genes associated with colon cancer linked single-nucleotide variants (SNVs) were utilized to create a protein-protein interaction (PPI) network. This was obtained from the MPRA screens for colon epithelial cells and HCT 116 cell line, each treated with butyrate and deoxycholic acid. The associated genes from both conditions were aggregated by cell type yielding two primary sets of genes for PPI network construction. These genes were queried against the STRING database, v12, species: Homo sapiens^83^. The query was restricted to interactions obtained through databases, experiments, coexpression, and text mining, with a high confidence (combined score ≥ 0.7). Genes that were resolved to the STRING database’s protein identifiers and had at least one interacting partner were used to construct the PPI network for downstream analyses. This network was further pruned to remove islands of binary interactors yielding the foundational networks for all further analyses. Network statistics such as node degree and edge betweenness were calculated and visualized using Cytoscape v3.10.3^84^. Node degree is a network property that indicates how many connections a node has, helping to identify proteins with numerous interaction partners that may function as key hubs. Edge betweenness, a complementary metric to node degree, quantifies the number of shortest paths passing through a particular edge, highlighting interactions that play essential roles in connecting different regions of the network. Additionally, Markov clustering^85^ was performed on the PPI network to identify modules in an unsupervised manner, which were then functionally enriched and multiple hypothesis testing correction was performed with a false discovery rate (FDR) limit of 0.05 (Benjamini–Hochberg procedure). The Gene Ontology database’s^86^ top two terms for Biological Processes were considered from the functional enrichment. For further annotation, proteins which interact with anti-neoplastic drugs were filtered from the Drug-Gene Interaction database v5.0.8^87^ and transcription factors (TFs) from the Human Transcription Factors database v1.01^88^.

### ChromBPNet Model Training

We trained ChromBPNet (https://github.com/kundajelab/ChromBPNet) models on bulk ATAC-seq profiles for each sample. For input, we used the peak and tagalign files for each sample, along with the pre-trained K562 bias model provided in the ChromBPNet repository (https://storage.googleapis.com/ChromBPNet_data/input_files/bias_models/ATAC/ENCSR868F GK_bias_fold_0.h5). We used a 5-fold cross-validation scheme to train the ChromBPNet models such that each chromosome appeared in the test set of at least one cross-validation fold. The chromosomes included in the train, validation, and test splits for each fold are the same as the ones from Marderstein et al. 2025 biorxiv (https://www.biorxiv.org/content/10.1101/2025.02.18.638922v2).

### ChromBPNet Variant Scoring

We used the ChromBPNet models for each sample to predict the base-resolution ATAC-seq coverage profiles for the 1 kb genomic sequence centered at each variant and containing the reference and alternate allele. Next, we estimated the variant’s effect size using two measures: (1) the log2 fold change in total predicted coverage (total counts) within each 1 kb window for the alternate versus reference allele, and (2) the Jensen–Shannon distance (JSD) between the base-resolution predicted probability profiles for the reference and alternate allele.

We determined statistical significance for both of these scores using empirical null distributions constructed by shuffling the 2114 bp sequence around each variant multiple times while preserving dinucleotide frequency. Each shuffled sequence was then duplicated and the variant’s reference or alternate allele was inserted at the center, resulting in a total of one million null variants for each set of observed variants scored. Each null variant was scored with the same procedure as the observed variants. For each observed variant, we computed the proportion of null variants with an equally high or higher (more extreme) score to derive empirical *P*-values for both the log2 fold change and JSD scores. The code base for scoring variants is at https://github.com/kundajelab/variant-scorer.

### LD-score regression analysis of colorectal cancer GWAS

Stratified LDSC analysis, as previously described by Finucane et al. (2018)^29^, was performed to obtain cell type-specific heritability estimates for stimulus-induced differentially accessible peaks and expressed genes using colorectal cancer GWAS summary statistics from the Lee Lab’s UKBiobank analysis^89^. Complete harmonized statistics for CRC GWAS were downloaded from the Lee Lab PheWeb database (https://www.leelabsg.org/resources). LDSC software was downloaded from Github (https://github.com/bulik/ldsc) and baseline model LD scores, weights, and allele frequencies (1000 Genomes European Phase 3) were downloaded from the Broad Institute Google Cloud storage platform (https://console.cloud.google.com/storage/browser/broad-alkesgroup-public-requester-pays/LDSCORE).

Stimulus-induced differential ATAC peaks were placed into the following buckets: 1) increased accessibility by butyrate (“Butyrate Open”); 2) decreased accessibility by butyrate (“Butyrate Closed”); 3) increased accessibility by DCA (“DCA Open”); 4) decreased accessibility by DCA (“DCA Closed”); 5) increased by butyrate AND decreased by DCA (“Butyrate Open & DCA Closed”); 6) decreased by butyrate AND increased by DCA (“Butyrate Closed & DCA Open”); 7) increased by both butyrate and DCA (“Butyrate Open & DCA Open”); 8) decreased by both butyrate and DCA (“Butyrate Closed & DCA Closed”).

Stimulus-induced differentially expressed genes were placed into the following buckets: 1) increased expression by butyrate (“Butyrate Hits Up”); 2) decreased expression by butyrate (“Butyrate Hits Down”); 3) increased expression by DCA (“DCA Hits Up”); 4) decreased expression by DCA (“DCA Hits Down”); 5) differentially expressed by butyrate in any direction (“Butyrate Hits All”); 6) differentially expressed by DCA in any direction (“DCA Hits All”); 7) significantly different between butyrate and DCA (“Butyrate v DCA Hits”).

Bucket-specific annotations were created from ATAC peaks and differentially expressed genes using the **make_annot.py** script. Background annotations for ATAC were separately created from the union set of all peaks. These annotations were added to the baseline model for LDSC heritability analysis to control for all open peaks and improve specificity of effects detected. Summary statistics were prepared for LDSC analyses using the **munge_sumstats.py** script with all default settings except chunksize=500000. To run cell type-specific heritability analysis, the LDSC script **ldsc.py** was run using the --h2-cts flag, with the background annotations added to the baseline model for ATAC (--ref-ld-chr) and testing for heritability enrichment in the bucket-specific annotations (--ref-ld-chr-cts).

## Notes

### Competing Interest Statement

The authors have declared no competing interest.

