## Supplemental Figures for "Interactions Between Dietary Metabolites and Regulatory Risk Variants for Human Colon Cancer"

#### Supplementary Figure 1

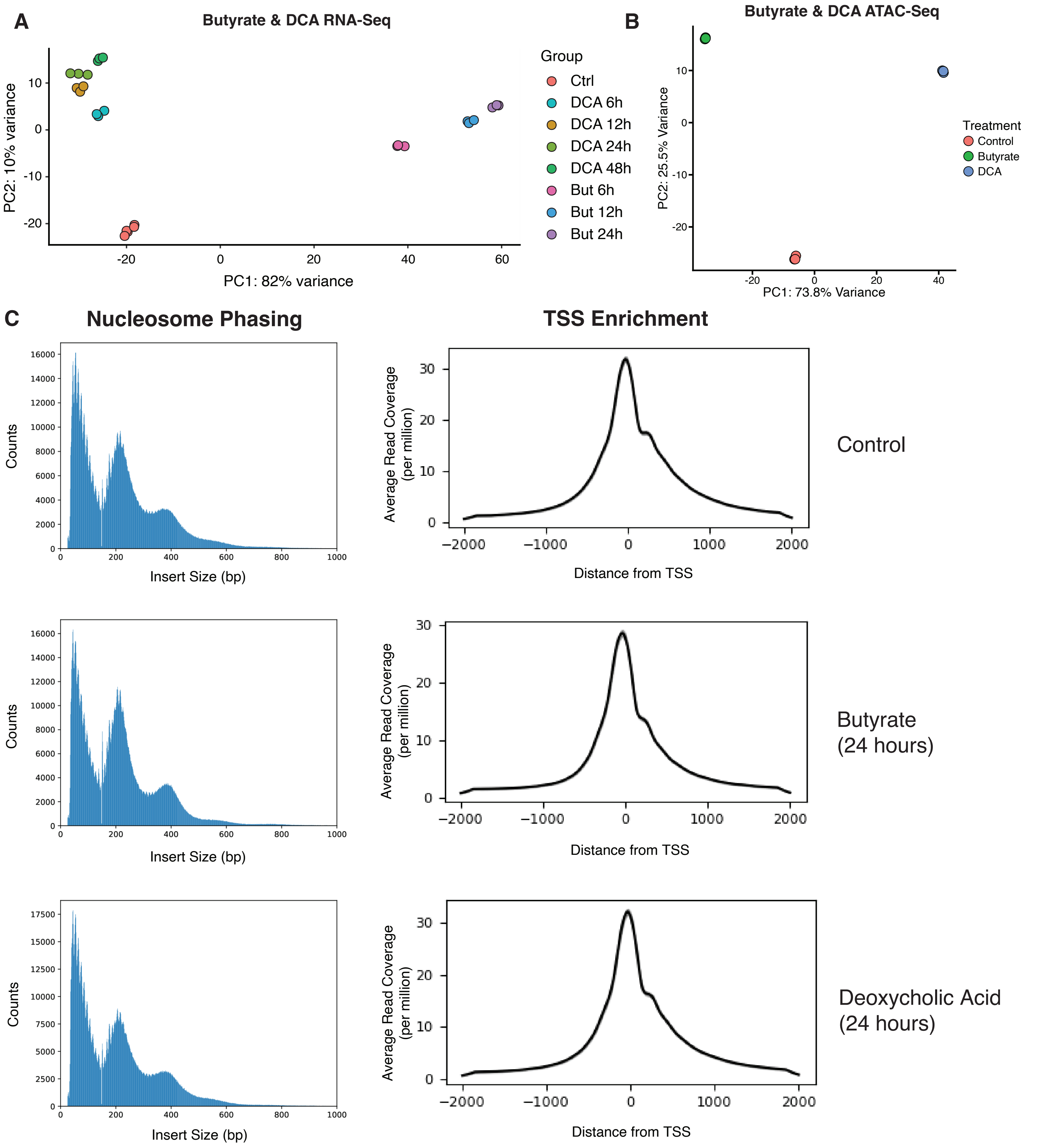

#### Supp. Figure 2

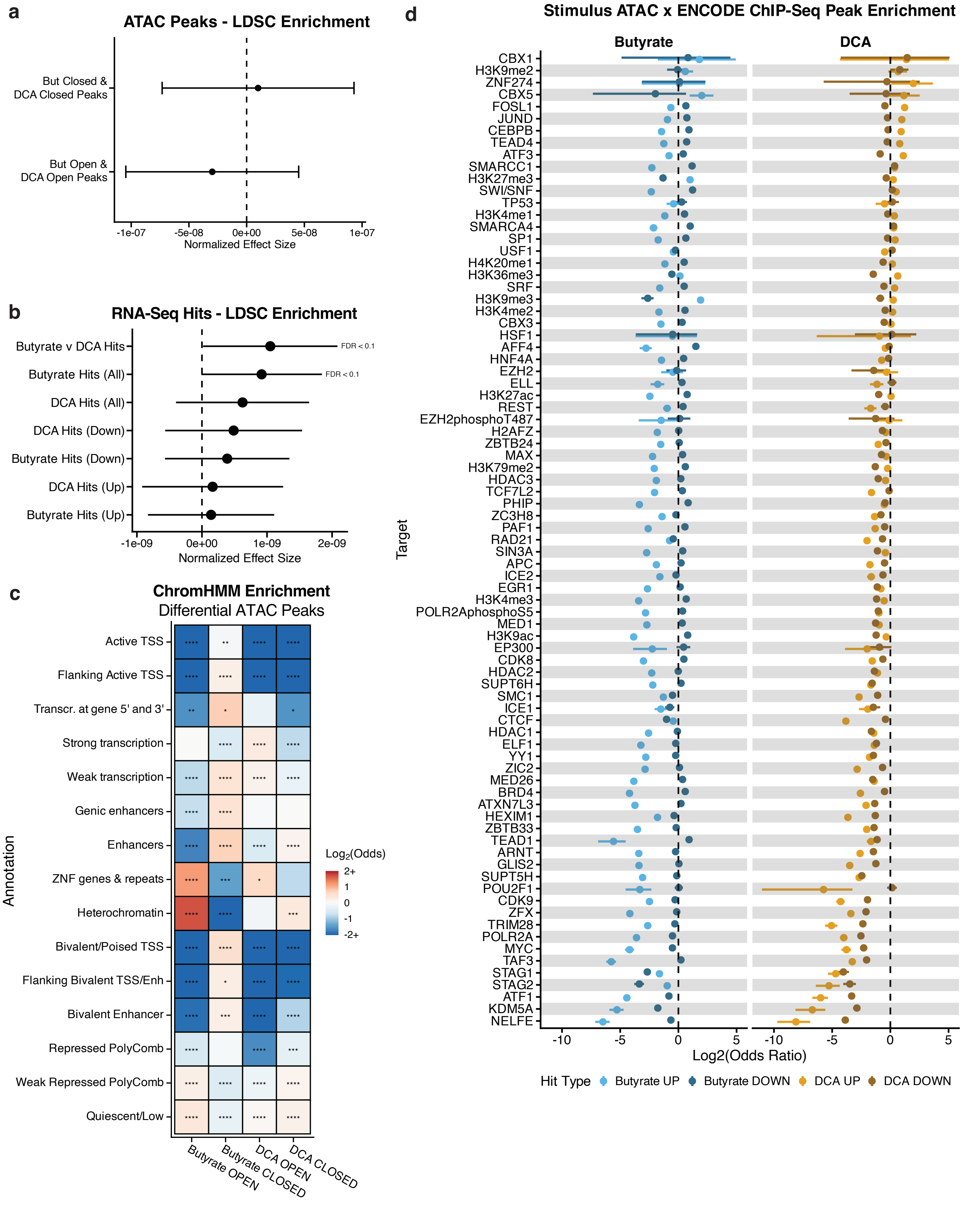

**Supp. Figure 3**

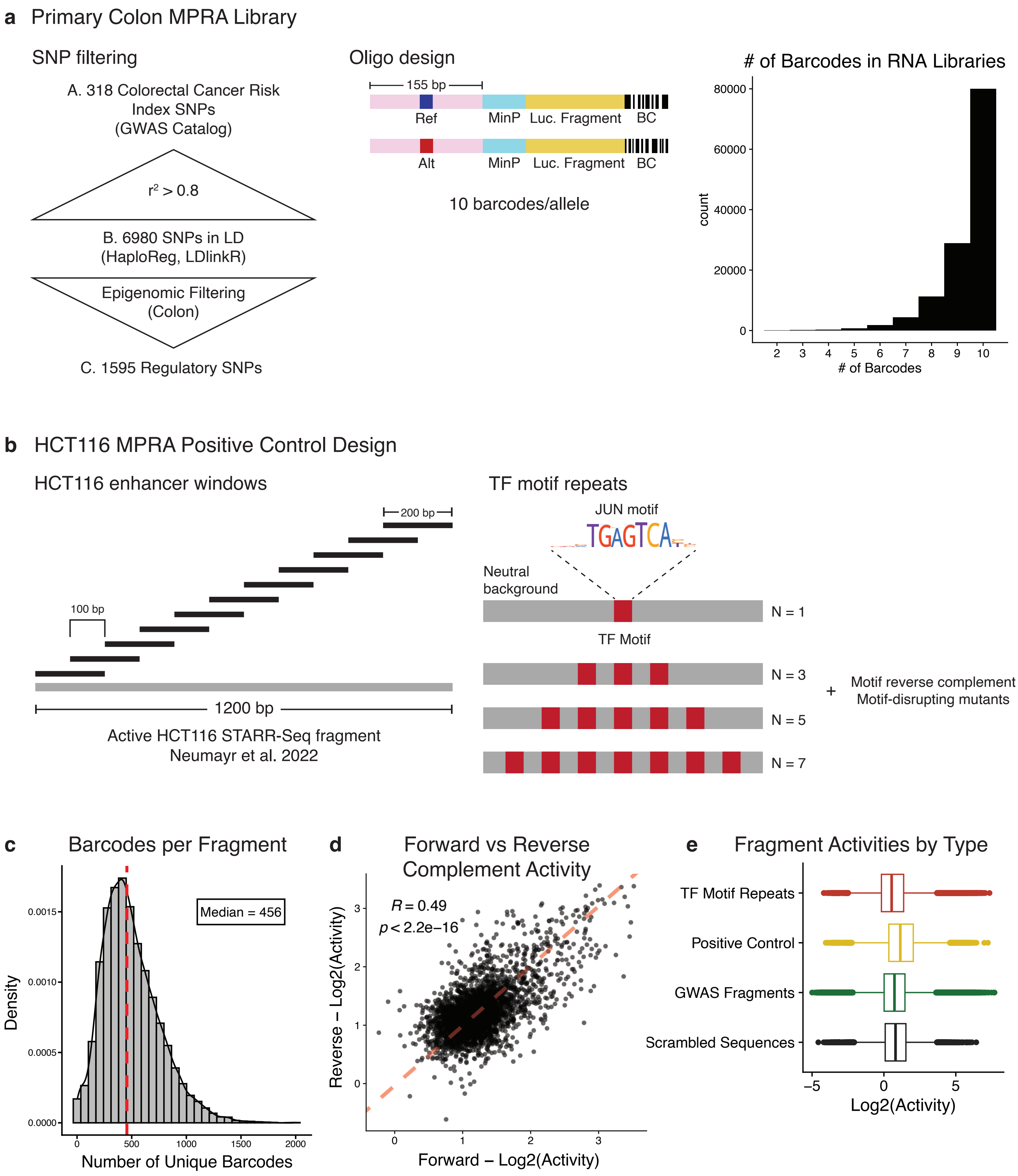

**Supp. Figure 4**

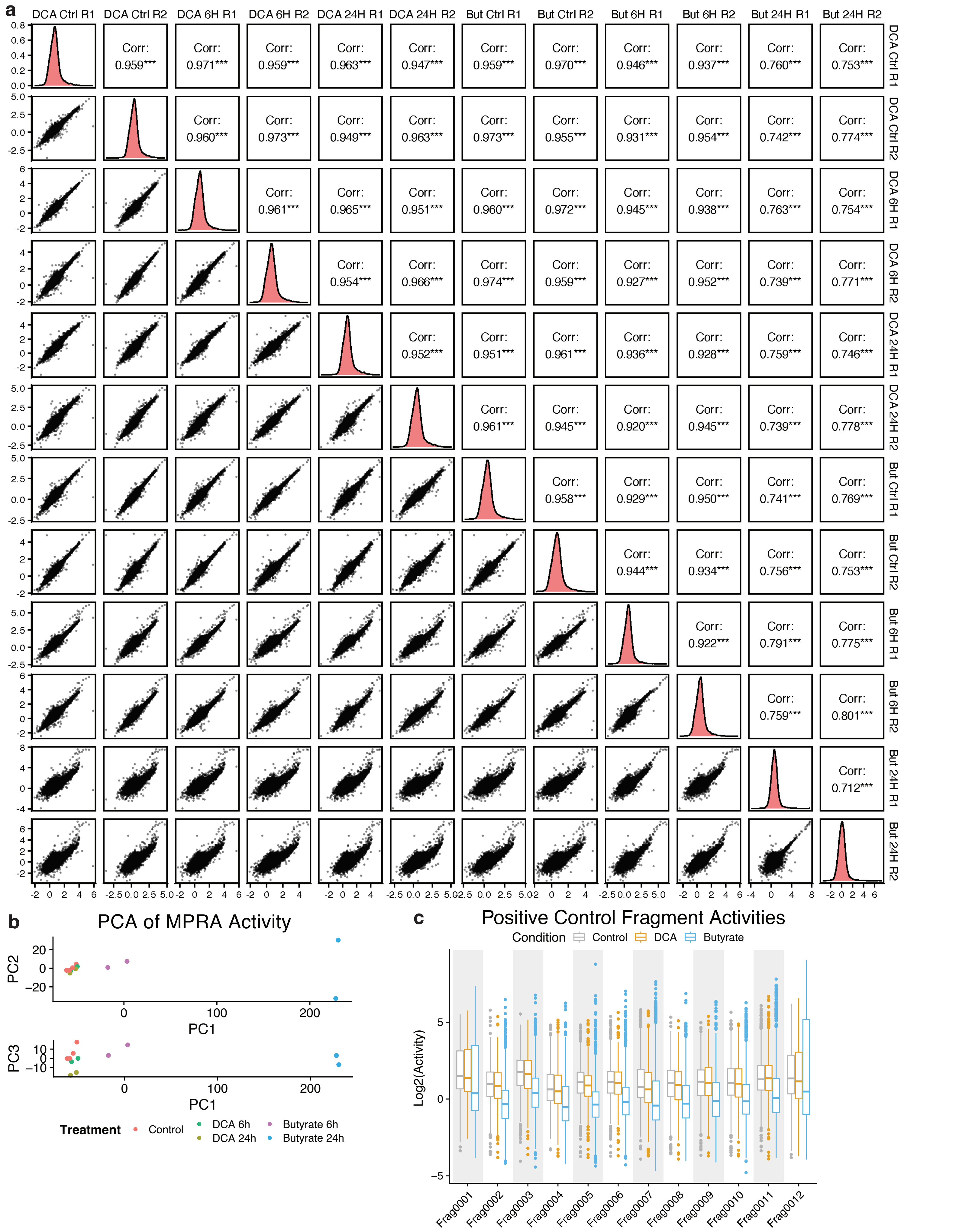

**Supp. Figure 5**

### TF Motif Repeat Activity by Condition

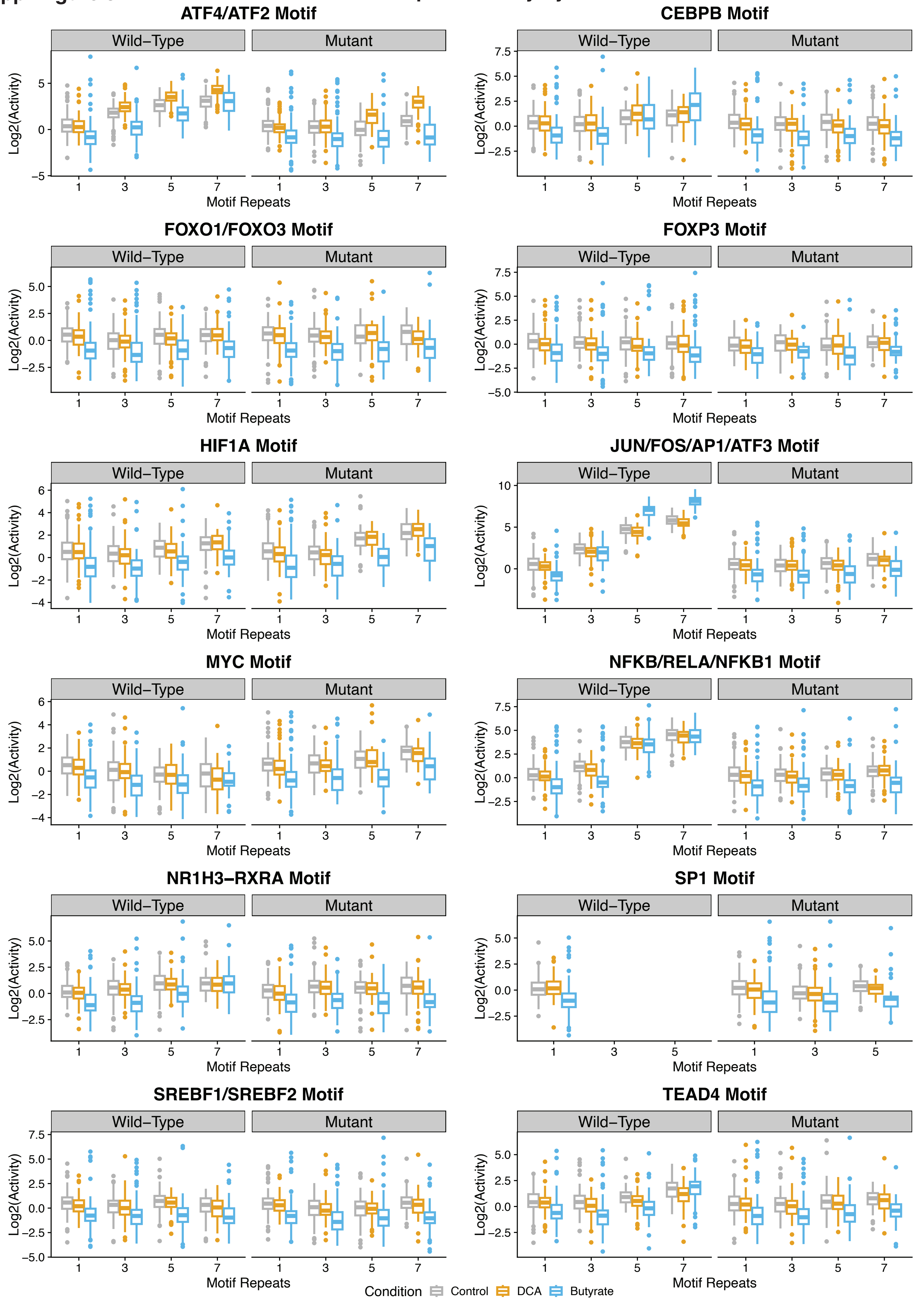

Supp. Figure 6

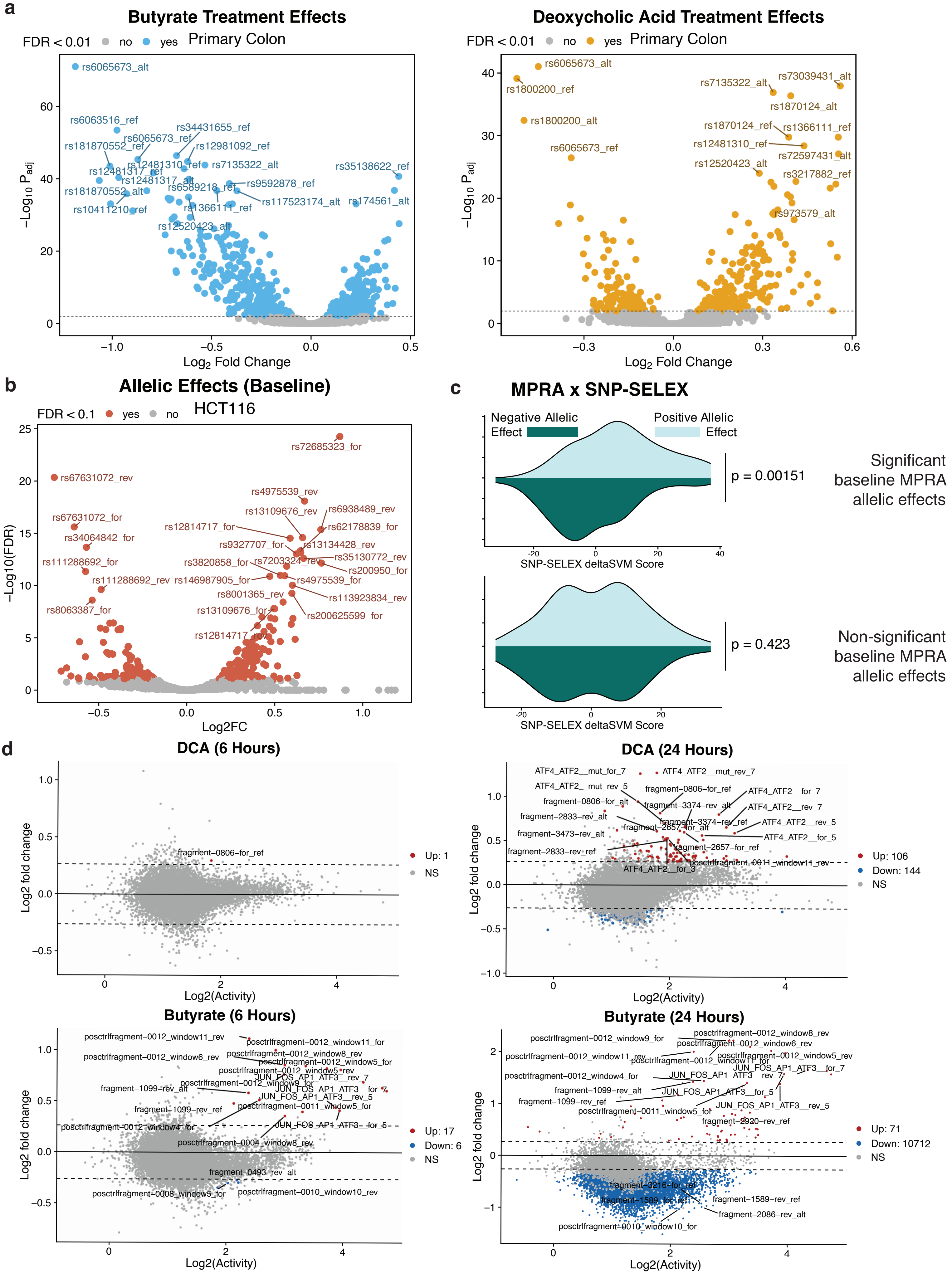

**Supp. Figure 7**

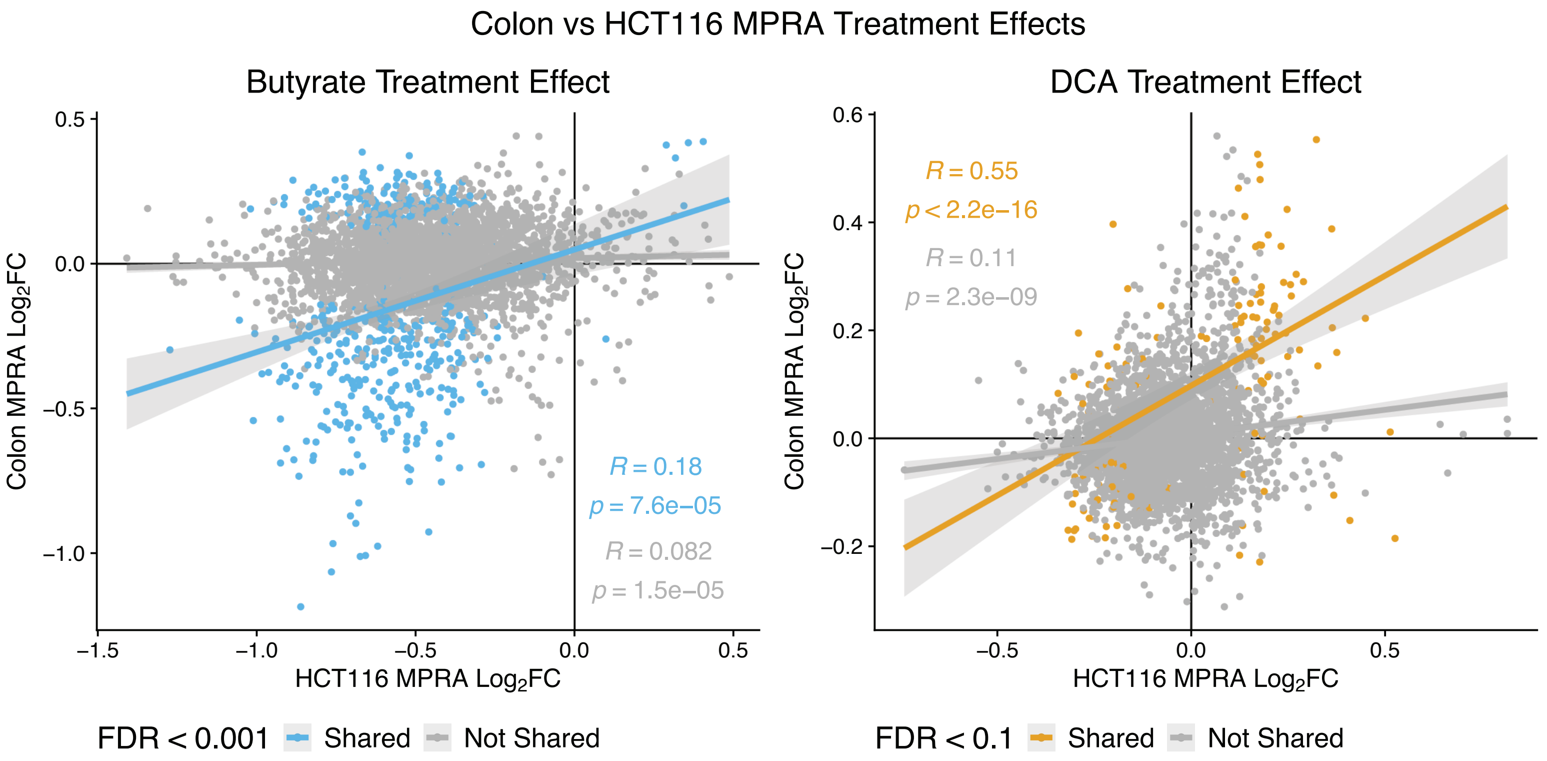

Supp. Figure 8

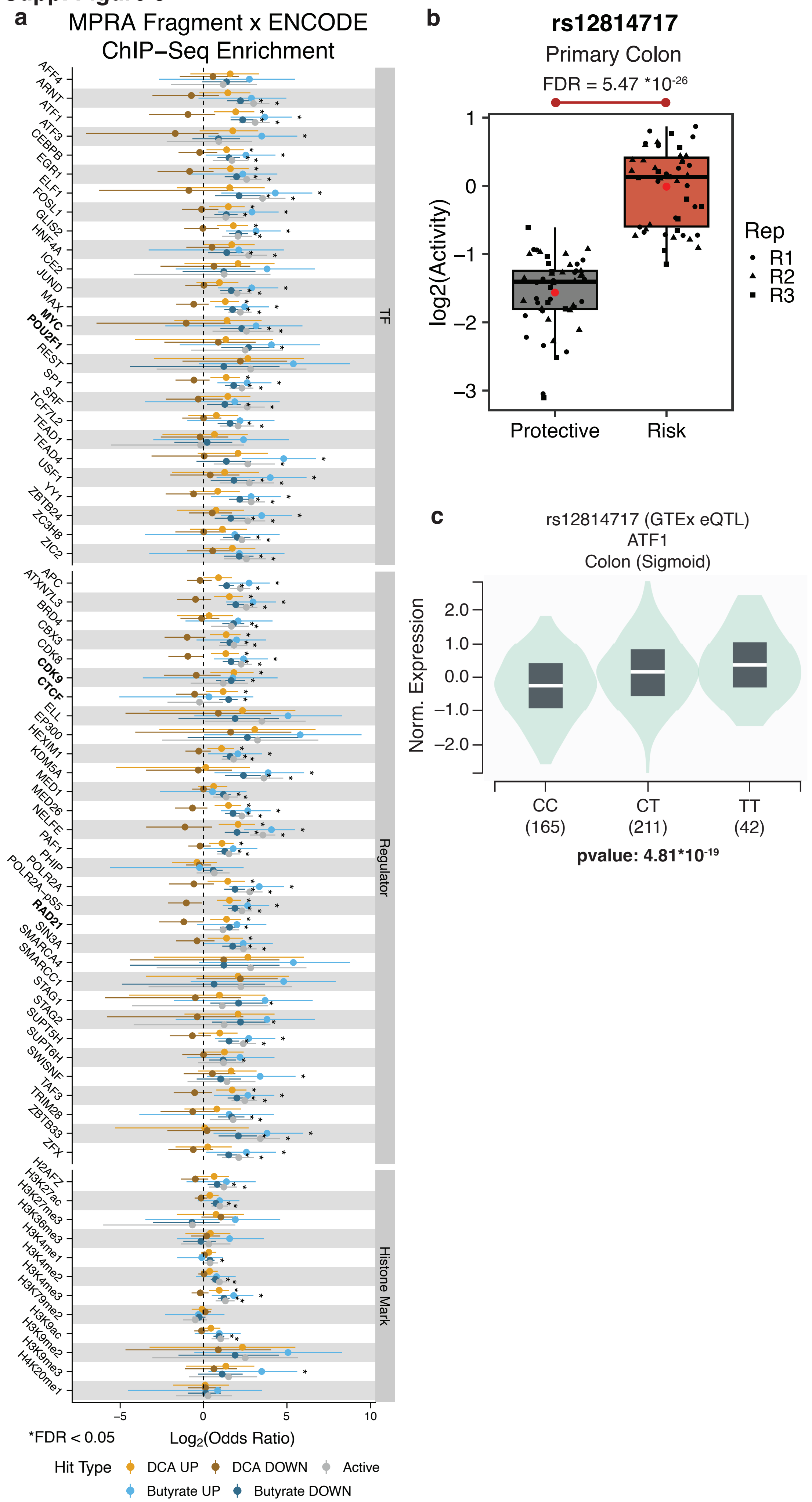

#### Supp. Figure 9

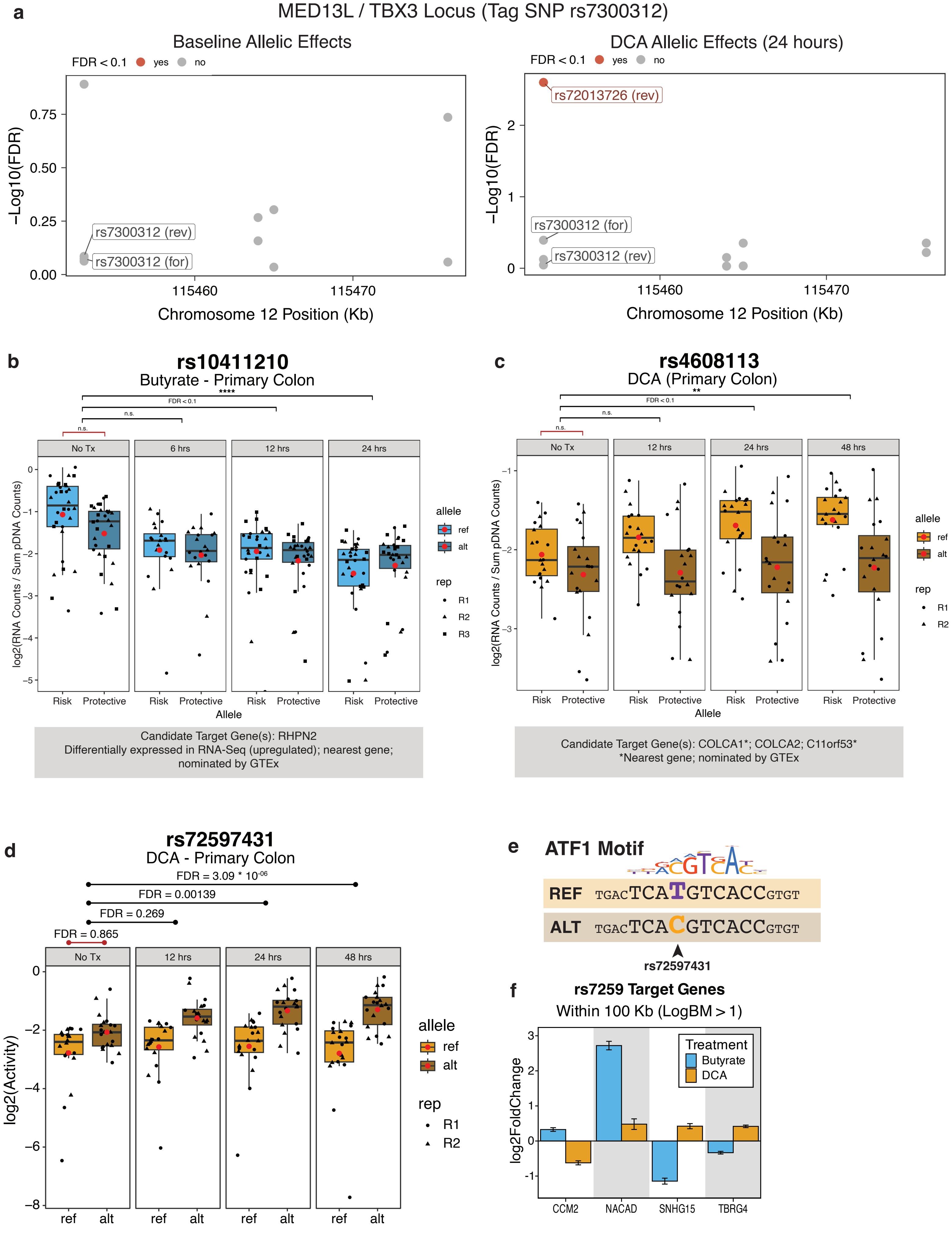

### Supp. Figure 10

a

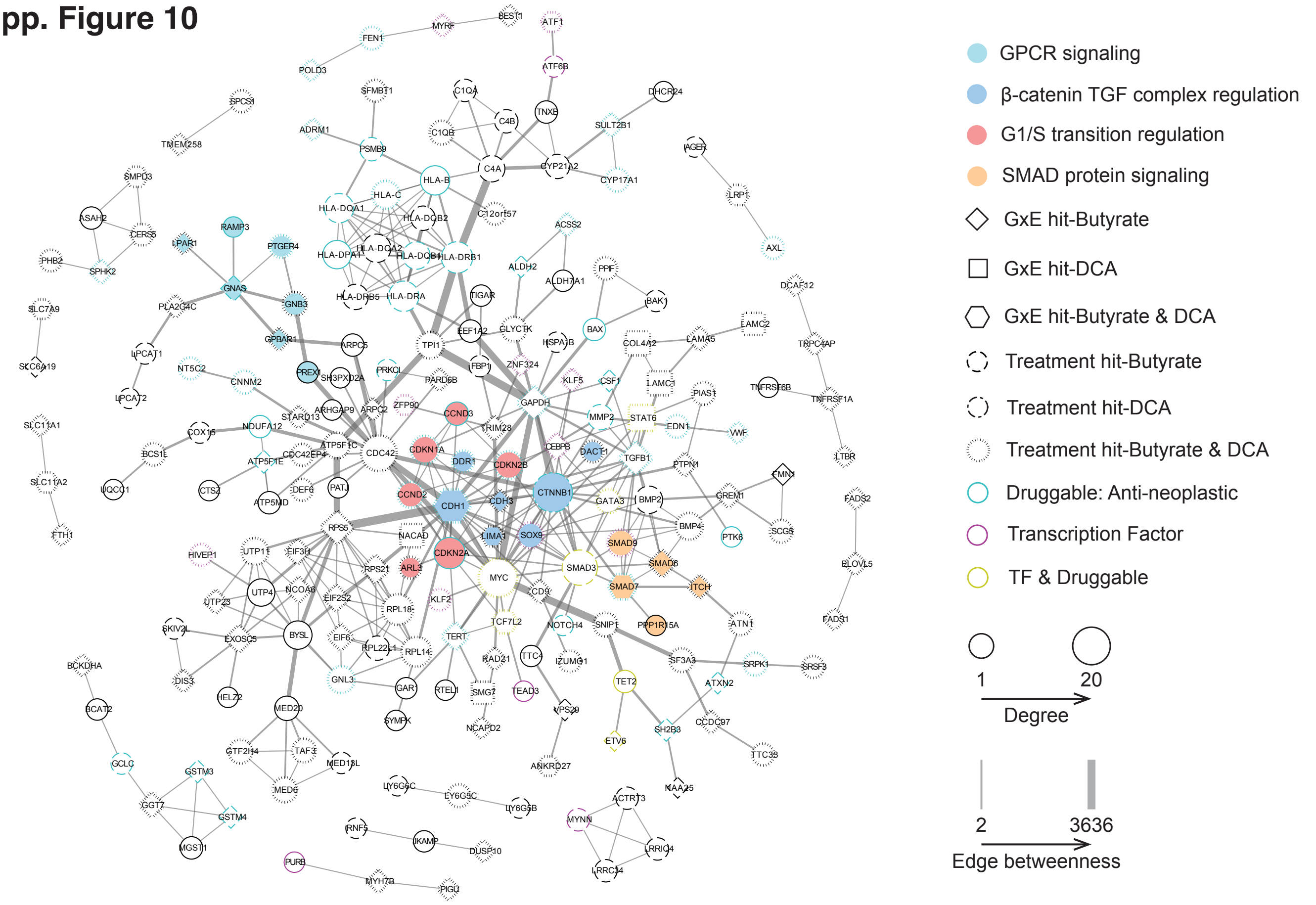


b

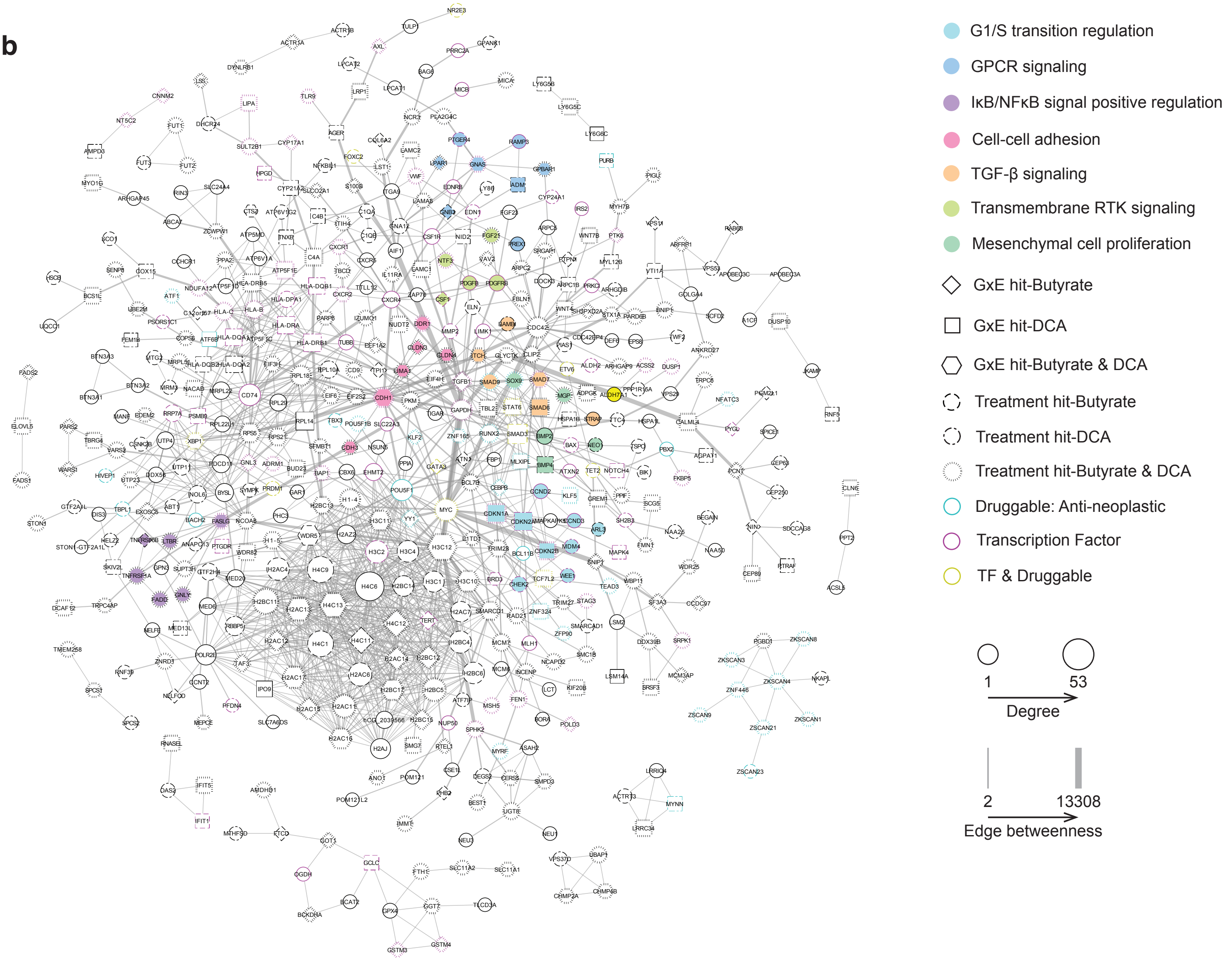
